# Detection of urease and carbonic anhydrase activity using a rapid and economical field test to assess microbially-induced carbonate precipitation

**DOI:** 10.1101/2020.01.10.902379

**Authors:** Fernando Medina Ferrer, Kathryn Hobart, Jake V. Bailey

## Abstract

Microbial precipitation of calcium carbonate has diverse engineering applications, from building and soil restoration, to carbon sequestration. Urease-mediated ureolysis and CO_2_ (de)hydration by carbonic anhydrase (CA) are known for their potential to precipitate carbonate minerals, yet many microbial community studies rely on marker gene or metagenomic approaches that are unable to determine *in situ* activity. Here, we developed fast and cost-effective tests for the field detection of urease and CA activity using pH-sensitive strips inside microcentrifuge tubes that change color in response to the reaction products of urease (NH_3_) and CA (CO_2_). Samples from a saline lake, a series of calcareous fens, and ferrous springs were assayed in the field, finding relatively high urease activity in lake samples, whereas CA activity was only detected in a ferrous spring. Incubations of lake microbes with urea resulted in significantly higher CaCO_3_ precipitation compared to incubations with a urease inhibitor. Therefore, the rapid assay indicated an on-site active metabolism potentially mediating carbonate mineralization. Field urease and CA activity assays complement molecular approaches and facilitate the search for carbonate-precipitating microbes and their *in situ* activity, which could be applied toward agriculture, engineering and carbon sequestration technologies.

## INTRODUCTION

Microbially-induced carbonate precipitation (MICP) has been explored as an alternative solution for several engineering and environmental challenges, such as restoration of building, monument, and concrete structures, soil consolidation, pollutant bioremediation and CO_2_ sequestration.^1–4^ Despite its relevance in the carbon cycle and its multiple applications, MICP is difficult to assess directly in a given environment. Current environmental microbiota analyses commonly characterize communities via 16S rRNA gene sequencing and metagenomic sequencing, which fail to determine active metabolisms, unless challenging and labor-intense transcriptomic or culture-based analysis are performed, which do not necessarily reflect *in situ* activity. It is possible, however, to directly test the activity of certain enzymes in the field and identify active metabolisms on-site. Here, we developed a method for the field detection of carbonic anhydrase (CA; EC 4.2.1.1) and urease (EC 3.5.1.5) activity, two enzymes associated with MICP.

Metabolisms such as photosynthesis, methane oxidation, nitrate reduction, bicarbonate transport and ureolysis can locally increase carbonate saturation and promote MICP.^4,5^ Two of these metabolisms rely on enzymes whose activity can be detected in the field: ureolysis and CO_2_ hydration via CA. Ureolysis, catalyzed by urease, allows microorganisms to use urea as a nitrogen and carbon source.^6^ CA facilitates rapid carbon transport into the cell via CO_2_ -HCO_3_^−^ interconversion.^7^ Both enzymes generate carbonate anions and increase the pH as a product of their activity. Urease generates ammonia and CO_2_ from urea (eq. 1), which coupled to ammonia hydrolysis and CO_2_ hydration, catalyzed by CA (eq. 2), produces one mol of hydroxide and one mol of bicarbonate per mol of urea (eq. 3):

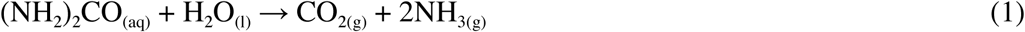

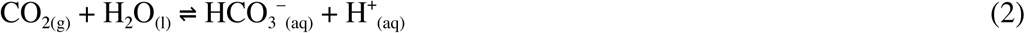

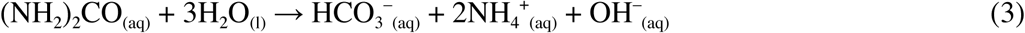

Urease and CA have been widely associated with MICP in a variety of environments.^4,8–10^ Ureolytic microbes have been applied in the restoration of buildings, soil consolidation and bioremediation,^3,11–15^ while CA has been proposed as an eco-friendly carbon sequestration technology to mitigate global warming.^2,16,17^ Yet, urease and CA effects are commonly evaluated via culture-based approaches, disregarding the activity of complex communities *in situ*. We evaluated on-site urease and CA activities using a rapid test applied to samples from calcareous fens, iron-rich springs, and a saline lake. In these settings, the environmental conditions may lead to carbonate precipitation, however, the contribution of MICP is difficult to assess in the absence of a field assay. By using a field test, we obtained a preliminary assessment of potential MICP, which can be applied to the management of urease- and CA-based technologies.

## MATERIALS AND METHODS

### Rapid urease field test

Urease activity was evaluated by a modification of the rapid urease test for *Helicobacter pylori* detection.^18,19^ The test consists of a sample-loading strip and an indicator strip (see a detailed preparation protocol in the supporting information). Sample-loading strips were prepared by immersing a cellulose paper in a fresh 0.6 M urea, 0.4 mM EDTA solution and drying the paper at room temperature. Indicator strips were similarly prepared by impregnating a paper with fresh 0.02% phenol red pH 6.0. Small strips of equal area (33 mm^2^) were cut from the urea-impregnated paper and circles of 6.5 mm diameter cut from the indicator-impregnated paper. Individual sample-loading strips were introduced in Seal-Rite 0.5 mL microcentrifuge tubes and indicator strips were placed inside the seals of the caps (**Figure 1A**). Urease activity was detected by adding standard solutions of urease from *Canavalia ensiformis* (Sigma-Aldrich, St. Louis, Missouri, USA) or wet biofilm samples in direct contact with the sample-loading strip before tightly closing the tube. A positive reaction was visualized by a color change from yellow to red in the indicator strip after ∼5 to 60 minutes of incubation at room/field temperature (**Figure 1B**). Volatile ammonia released from urea (eq. 1) increases the pH in the indicator strip, changing its color (**Figure 1C**). A negative control was prepared under the same conditions and additionally adding 1 mM phenyl phosphorodiamidate (PPD)—a urease inhibitor—in the soaking solution of sample-loading strips. A positive reaction without the inhibitor and a negative reaction with PPD strongly indicate urease activity.

**Figure 1.**
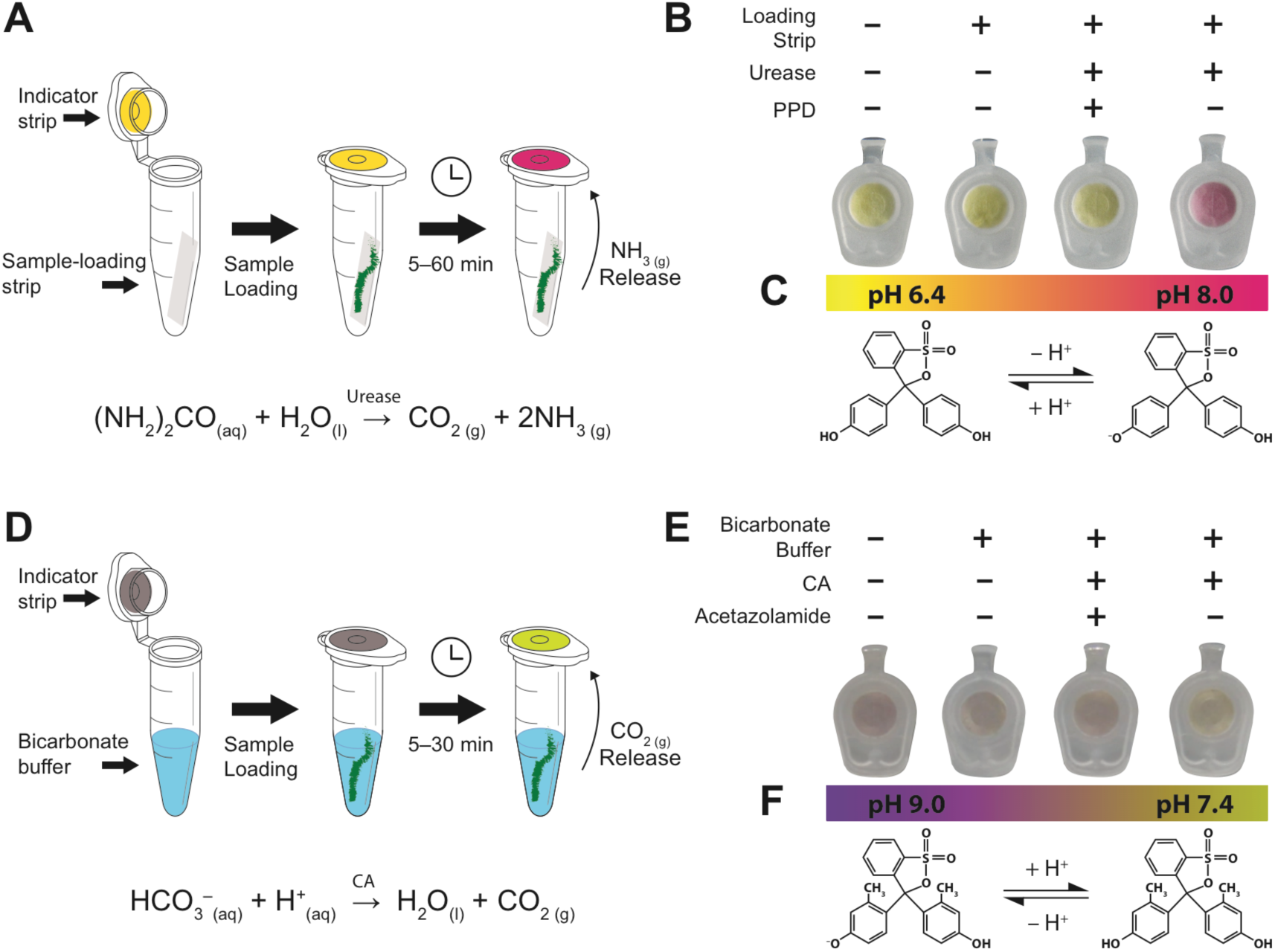
Urease and CA field tests. (A) Representation of a positive urease assay after adding a biofilm sample in direct contact with the loading strip. (B) A switch to red in the tube cap (indicator strip) indicates a positive urease reaction. If no urease is present or if a loading strip with urease inhibitor (PPD) is assayed, the indicator is expected to remain yellow. (C) Color transition of phenol red (pKa 7.4, pH range 6.4–8.0) used for urease tests. (D) CA assay scheme after adding a sample in the tube with bicarbonate solution. (E) Faster development of a yellow color in the indicator than a negative control (no sample added or using a CA inhibitor) indicates positive CA activity. (F) Color transition of metacresol purple (pKa 8.32, pH range 7.4–9.0) used for CA assays.

### Rapid CA field test

CA assays were prepared using a CO_2_ detection method intended for proper endotracheal catheter introduction.^20^ CO_2_-sensitive strips were prepared by soaking a cellulose paper with fresh 0.0065 M Na_2_CO_3_, 0.01% metacresol purple, 50% glycerol diluted with N_2_-purged distilled water (see detailed preparation protocol in the supporting information). Impregnated papers were immediately dried by a stream of hot air and circles of 6.5 mm were cut and placed inside the cap seals of 0.5 mL tubes (**Figure 1D**). The bluish-purple indicator gradually turns purplish-yellow after one to three days of exposure to atmospheric CO_2_. We either used freshly prepared indicator strips or stored them for few days inside a tube containing Ca(OH)_2_ to minimize color changes. CA activity was detected by introducing 80 μL of cold 1 M NaHCO_3_ in the tubes and adding the samples or 15 μL of standard CA (isozyme II from bovine erythrocytes; Sigma-Aldrich, St. Louis, Missouri, USA) solutions. The tubes were immediately closed and incubated on ice. Bicarbonate dehydration (catalyzed by CA, eq. 2) produces volatile CO_2_, which reacts with glycerol-absorbed water in the indicator lid and generates acidity that turns the indicator from bluish-purple to purplish-yellow (**Figure 1E**). Non-enzymatic bicarbonate dehydration proceeds rapidly and therefore the indicator color change is observed within minutes, even without the enzyme. By using the CO_2_-sensitive strips in microcentrifuge tubes we found a subtle, but reproducible, color change difference between CA-incubated (10–100 mM CA) and negative controls. Negative controls were prepared under the same conditions and adding 5 μL of fresh 1 mM acetazolamide—a CA inhibitor—to the bicarbonate buffer. A faster color change compared to negative controls is indicative of CA activity.

### Microbial sampling locations

Samples were taken near calcareous fens in the Minnesota River Basin, from ferrous springs and from Salt Lake, MN, during July and August 2019 (see detailed locations in **Figure S1** and **Table S1)**. Salt Lake is an alkaline sulfate- and sodium-dominated saline lake.^21^ The lake alkalinity (234 ± 2 mg/L CaCO3, pH 8–9), together with calcium carbonates in its sediments,^21^ indicates favorable conditions for carbonate mineral precipitation, which could be stimulated by microbial metabolisms. A sample of buoyant green biomass was collected from Salt Lake during a bloom event in July 2019. Submerged green filaments attached to shoreland rushes were also collected following the bloom in August 2019 (**Figure S2**).

Calcareous fens, peatlands in which surficial calcium carbonate precipitates,^22^ are also environments where microorganisms may contribute to carbonate mineralization. Samples were collected from green biofilms growing on peat exposed by a creek at Black Dog Lake Fen, and from surficial green filaments suspended on water ponds during flood events in Nicols Meadow Fen, and between Fontier 8 Fen and Sioux Nation WMA Fen (locations shown in **Table S1** and **Figure S3**). Additionally, samples from iron-oxidizing microbes were collected from orange precipitates at a creek in Nicols Meadow Fen, from groundwater seeping to the Mississippi River at Saint Mary’s Spring, and from a sulfide seep at a roadside near Soudan, MN (**Figure S4**). A small portion of biomass (enough to wet the sample-loading strip of the urease assay) was evaluated on-site in triplicate (with the exception of the Salt Lake bloom sample, where only two samples were evaluated) for urease and CA activity using the rapid field test. Additional triplicate aliquots for each sample were obtained for protein extraction and microscopic observations (supporting information).

### Calcium depletion kinetics

Salt Lake filaments (Sample ID 02 in **Table S1**) were incubated (0.5 g wet weight) in 20 mL of 0.2 μm-filtered lake water with the addition of 0.8 M urea in closed 50-mL glass serum bottles with agitation (90 rpm, orbital shaker MaxQTM 2000, Thermo Fisher Scientific) at room temperature and under natural light cycles for 12 days. Four different incubation conditions were evaluated in triplicate: lake water without inhibitors, lake water with 1 mM acetazolamide, lake water with 1 mM PPD, and lake water with both inhibitors (1 mM each). Aliquots of 0.5 mL were taken over time and titrated with EDTA (HAC-DT, Hach, Loveland, CO, USA) to quantify calcium. Solid residues after incubations were evaluated by powder micro X-ray diffraction (micro-XRD) using a Bruker D8 Discover micro-diffractometer with a CoKα source (λ = 1.78899 Å) equipped with a graphite monochromator and a 2D Vantec 500 detector. Samples were mounted on vacuum grease and three frames (30° 2θ width) centered at 20°, 45° and 70° were collected for 900 s at 40 kV and 35 mA. Phase identification was conducted using Match! (v3.8.3.151) and the Crystallographic Open Database (COD-Inorg REV218120 2019.09.10) reference patterns for aragonite (96-901-6601), calcite (96-900-0971), monohydrocalcite (96-901-2074), quartz (96-901-0145), thenardite (96-900-4093) and vaterite (96-150-8972). Ikaite diffraction pattern was obtained from Hesse and Kueppers (1983).^23^

### Statistical analysis

To compare and semi-quantify the rapid test results, we followed color changes using hue values. The hue represents color pigmentation by a single number, disregarding saturation and brightness, therefore, minimizing color differences resulting from light and exposure time changes in the field (**Figure S5**). Average hue values were calculated from RGB colors of standard circle areas over photographs of the indicator strips using the NIH ImageJ 1.49v software. Statistical significance in the rapid assays and in the calcium depletion kinetics was assessed via a Student’s t-test using GraphPad Prism 5.0.

## RESULTS AND DISCUSSION

### Rapid test sensitivity and reproducibility

A pH-sensitive dye encapsulated within a cellulose matrix was used as a detector of NH_3_ or CO_2_ inside the cap of microcentrifuge tubes. Urease activity was followed by an ammonia-mediated pH increase, whereas CA was detected by a pH decrease. The test format in **Figure 1** turns a phenol red indicator from yellow to red in less than 15 minutes when >30 mU urease is assayed (**Figure 2A**). After 30 minutes, the method sensitivity is ∼3 mU, when compared to a PPD-containing negative control (**Figure 2B**).

**Figure 2.**
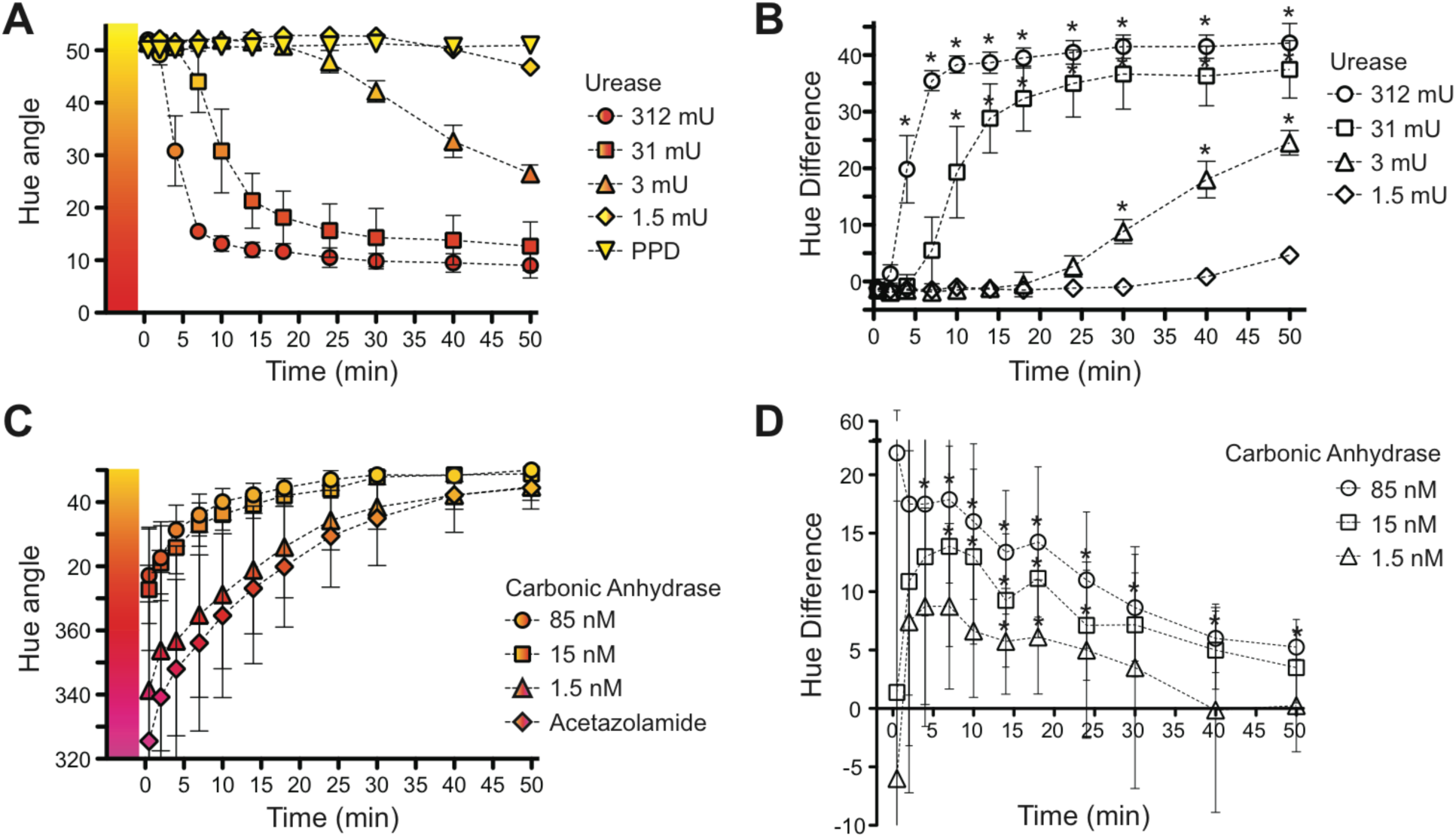
Field test color change kinetics and sensitivities. (A) Assay color (expressed as average hue values) as a function of incubation time for the urease test using different urease standards (1.5–312 mU), and the average of their negative controls containing a urease inhibitor (PPD). (B) Hue difference of urease assays compared to their negative controls. (C) CA assay hue as a function of incubation time after adding different CA concentrations (1.5–85 nM), and the average of their negative controls in the presence of a CA inhibitor (acetazolamide). (D) Hue difference of CA standards compared to their negative controls. All conditions were assayed using six replicates. Error bars represent standard deviations. *p < 0.05 between each condition and its respective negative control.

For CA assays, however, the rapid and spontaneous bicarbonate dehydration, even without the enzyme, turns metacresol purple indicator to yellow within 30 minutes (**Figure 2C**). The reaction has a window of 20–30 minutes where a coloration difference is noticeable between a CA-containing assay and an acetazolamide-containing test (negative control), with a maximum hue difference observed between 2 and 20 min (**Figure 2D**). When using 1.5 nM CA, we observed a subtle, but consistent, hue difference that was significant at 14 and 18 minutes.

### Field detection of urease and CA activity

Urease and CA tests were used for on-site enzymatic activity detection in samples from a saline lake, ponds from calcareous fens, and iron oxide precipitates from ferrous springs. Biomass collected during a bloom event in Salt Lake and a sample of green filaments collected after the bloom were urease-positive in less than 20 minutes (**Figure 3A**).

**Figure 3.**
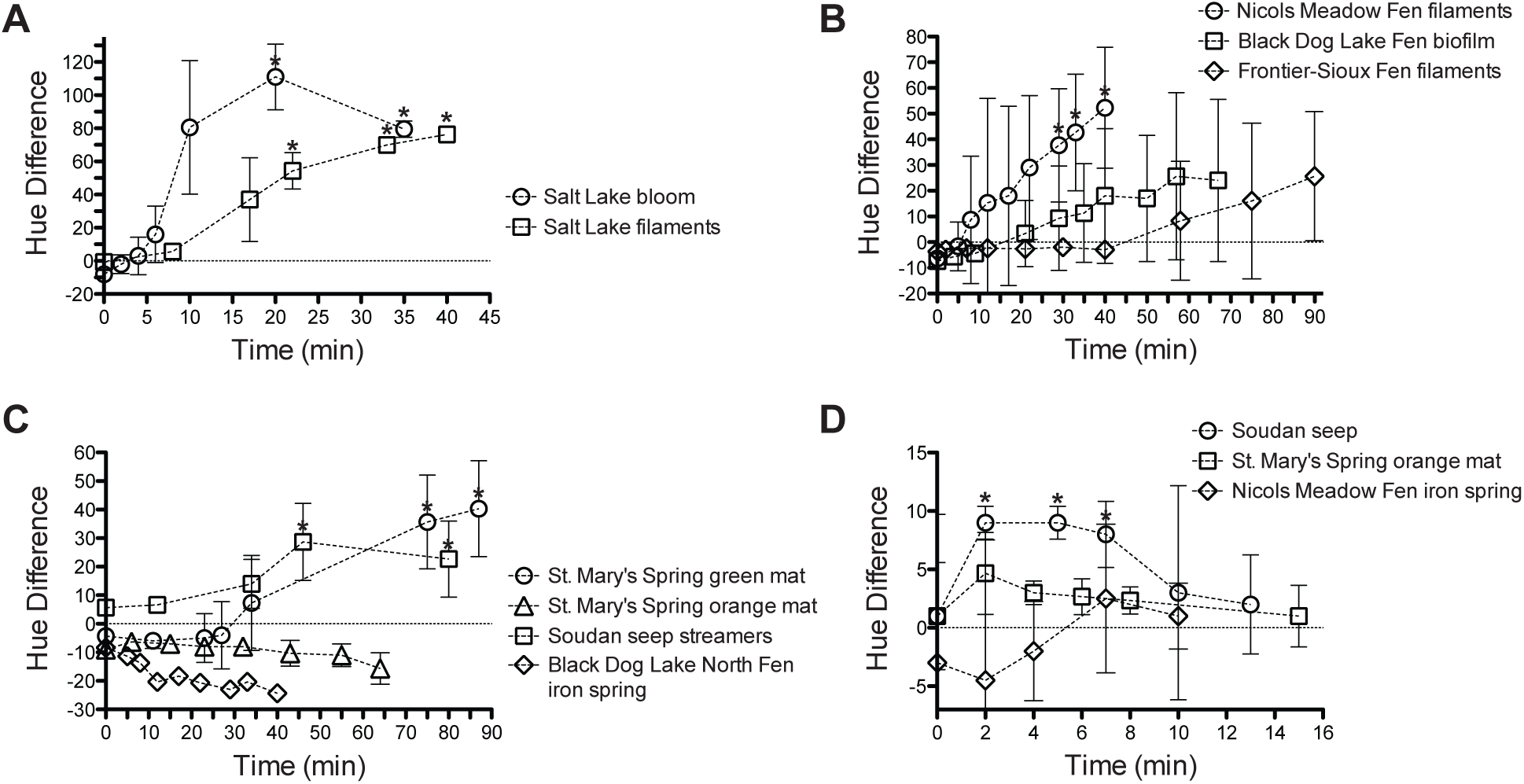
Field test color change using environmental samples. Hue difference from on-site urease assays of Salt Lake (A), calcareous fen (B), and iron spring (C) samples. (D) CA assay hue difference of ferric precipitates. Error bars represent standard deviations. *p < 0.05 between each sample and its respective negative control.

Black Dog Lake Fen and Nicols Meadow Fen samples were also positive for urease, although longer incubations were required and they showed less color change intensity than the samples from Salt Lake (**Figure 3B**). Green photosynthetic sheaths from Frontier-Sioux Nation Fen showed little urease activity, even after one hour of incubation. In organisms that do not constitutively express urease, its expression is likely induced when urea is available.^24,25^ The higher urease activity in Salt Lake may therefore reflect urea accessibility and correlate with the bloom event observed in July 2019. Agriculture promotes eutrophication,^26^ and in particular urea—the major worldwide fertilizer^27^—can be used by microbes as both N and C source.^6^ Salt Lake is located in close proximity to farmland and its microbial communities may be sensitive to nearby fertilization practices. By contrast, less urease activity found in calcareous fen samples, in particular Frontier-Sioux Nation Fen, which is located near a State Wildlife Management Area, may reflect a lower agriculture impact (**Figure S6**).

Biofilms at St. Mary’s Spring have a combination of green cyanobacterial filaments that were slightly positive for urease, and stalks of iron-oxidizing bacteria surrounded by few cyanobacteria and iron oxides, which were urease-negative (**Figure 3C**). Iron oxides from Black Dog Lake North Fen creek were also urease-negative. PPD-containing controls in these ferric precipitates, however, showed a slight color change (represented by a negative hue difference in **Figure 3C**), attributed to ammonia release from PPD degradation. The P–N bonds in phosphoramidates are unstable in aqueous solutions^28^ and may release ammonia. Moderate transitions to red may lead to false positives, although its intensity was not comparable to the positive reactions observed in other environments. Several other urease inhibitors not tested in this study^29^ may prevent false positives, however, phosphoramidates (such as PPD) are among the most potent and specific urease inhibitors^28^ and were therefore selected for the assay.

In contrast to urease, CA activity was not detected in Salt Lake and calcareous fen samples. Only Soudan seep samples were positive for CA when comparing hue values with those of negative controls (**Figure 3D**). CA is essential for carbon transport and pH regulation.^7^ Microbes from Soudan sulfidic seeps were located at a site where recent road construction exposed sulfide outcrops that potentially generate acid rock drainage. As a mitigation attempt implemented by the Minnesota Department of Transportation, limestone was placed at the roadsides, affecting microbial populations that likely overexpress CA to tolerate high alkalinity (158 ± 4 mg/L CaCO_3_) and pH fluctuations. CA is also fundamental to CO_2_ concentrating mechanisms in photosynthesis; therefore, its expression is expected in photosynthetic biofilms.^7^ It is possible that CA levels were below the detection limit of the field assay. While the urease assay was shown to be robust and shown to have low detection limits, the CA assay was limited by the nature of its catalyzed reaction (eq. 2), where only few minutes are available to visualize CA activity. In addition, carbonic anhydrases include at least six enzyme classes with no structural or sequence homology and, consequently, inhibitors may have varied effects. Although acetazolamide is a potent wide-spectrum CA inhibitor,^30,31^ it is possible that certain microbial CA were not effectively inhibited, hindering a positive reaction. Alternatively, the enzyme may not have been readily accessible to the bicarbonate substrate and consequently the non-enzymatic reaction masked CA activity. Intracellular CA requires bicarbonate transport to the interior of the cell^32^ and therefore the color change detection is dependent on CO_2_ escape from the cell interior. Additional transport processes may delay CO_2_ generation in the reaction tube. We failed, however, to obtain environmental protein extracts with significant soluble CA activity (**Figure S7**), even in photosynthetic cells. We also found very low soluble urease activity from protein extracts (**Figure S7**), indicating that most on-site activity found may have been the result of extracellular, possibly membrane bound or extracellular polymeric substances (EPS)-associated urease, which was observed as residual urease activity of cell debris after protein extraction.

### Carbonate precipitation induced by urease and CA

Differences in intensity and time to obtain a positive reaction serve as parameters to compare relative activities among sites, where the strongest urease activity was found in Salt Lake samples. To determine the potential MICP of urease and CA, we incubated Salt Lake filaments with lake water containing urea, showing a decrease in its Ca^2+^ concentration (14.4 ± 0.5 mM) starting at Day 4, and depleting at Day 9 (**Figure 4A**). The calcium drop is interpreted as calcium carbonate precipitation, which was also observed by the solution turbidity (attributable to CaCO_3_) starting at Day 3–4 (**Figure 4B–D**). By contrast, incubations in the presence of PPD depleted only around one-third of the initial calcium after 12 days, having significantly higher calcium than incubations without PPD after Day 4 (**Figure 4A**).

**Figure 4.**
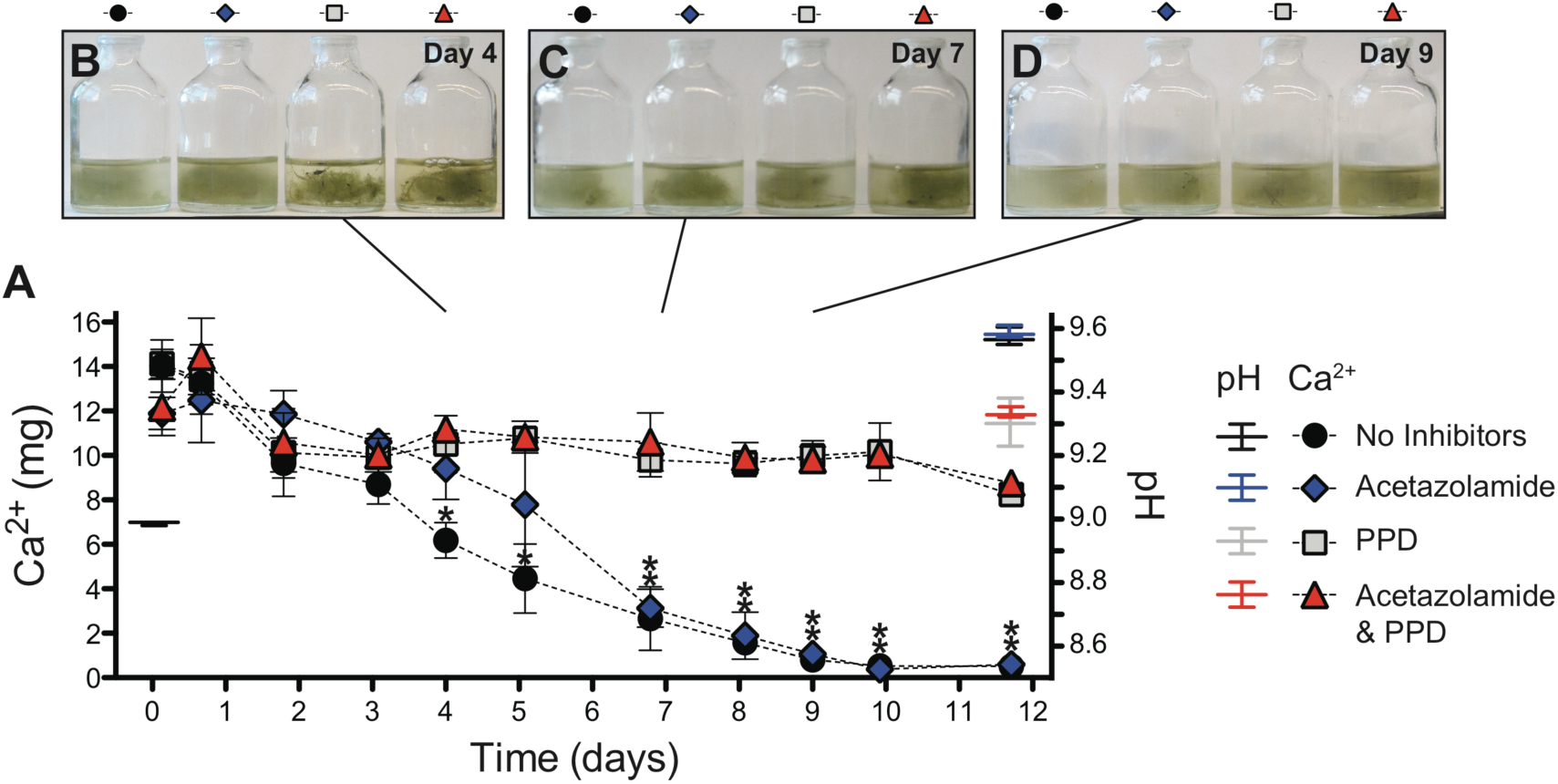
Calcium depletion kinetics of Salt Lake water in the presence of its microbial biomass. (A) Calcium quantification (left axis) from aliquots taken over 12 days for triplicate incubations with a CA inhibitor (acetazolamide), a urease inhibitor (PPD) and conditions with and without both inhibitors. A pH increase (right axis) is observed at the end of the incubations. Standard deviation of triplicate assays are represented by the error bars, *p < 0.05. Photos of representative incubations at Day 4 (B), 7 (C) and 9 (D) show hazy solutions attributable to mineral precipitation.

Filament incubations in the presence of acetazolamide, however, did not prevent calcium depletion (**Figure 4**). Therefore, urease, and not CA activity, is most likely responsible for carbonate precipitation under the studied conditions, consistent with the enzymatic activity observed in the field. Between Days 4 and 7, however, a delay in calcium depletion with acetazolamide compared to the condition without inhibitor may indicate a possible CA influence on CaCO_3_ precipitation dynamics. Both enzymes may synergistically affect carbon precipitation, as suggested previously,^33^ via rapid bicarbonate generation by CA (eq. 2) from CO_2_ produced by urease (eq. 1), providing CO_3_^2−^ and neutralizing the acidity produced by CO_2_ hydration. The mechanism by which urease promoted CaCO_3_ precipitation is attributable to a pH increase during incubations compared to the lake water initial pH (**Figure 4A**). Without PPD, the pH rises more than 0.5 units in 12 days, increasing the saturation with respect to carbonate minerals (**Figure S8**).

Following 12 days, the filaments were covered by a white precipitate, extensively found in incubations without inhibitors or with acetazolamide only. Under the microscope, green trichomes were encrusted by precipitates, which appeared white under phase contrast, blackish under bright field and were autofluorescent (**Figure 5A**). The encrusting particles may correspond to magnesium-containing calcium carbonates, which have been observed to emit wide-spectrum fluorescence.^34^ Additionally, characteristic signals of calcium carbonate polymorphs, such as vaterite, monohydrocalcite, calcite, ikaite, and aragonite, along with thenardite (Na_2_SO_4_, likely the result of high Na and SO_4_ in Salt Lake) and quartz (presumably from diatom frustules) were detected in the precipitates by micro-XRD (**Figure 5B**).

**Figure 5.**
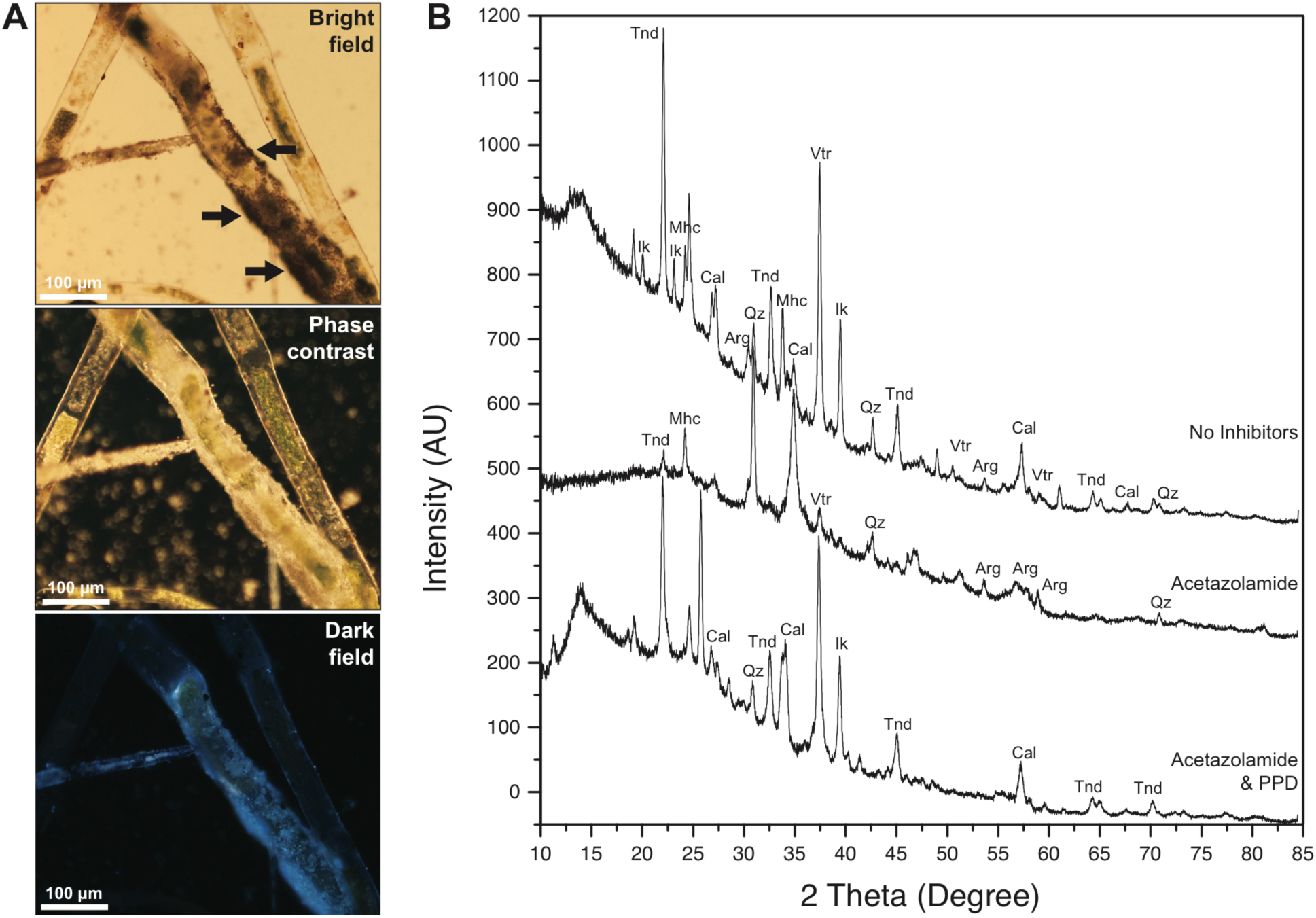
Salt Lake microbial sheaths after incubations. (A) Filament photomicrographs after 12 days of incubation, showing black precipitates covering the sheaths under bright field (black arrows, top panel) that appeared white under phase contrast (middle panel) and have wide-spectrum autofluorescence (lower panel, dark field showing 420 nm-excited 450 nm long-pass emission). (B) Micro-XRD of precipitates after incubations without inhibitors, with acetazolamide, and with both inhibitors. Mineral abbreviations: Arg, aragonite; Cal, calcite; Ik, ikaite; Mhc, monohydrocalcite; Qz, quartz; Tnd, thenardite; Vtr, vaterite.

### Potential applications of urease and CA field tests

The field detection of urease activity could prove useful for evaluating the predictability of fertilizer efficiency. Urea-based fertilizers are hydrolyzed by soil bacteria, resulting in volatilization and nitrogen loss to the atmosphere, which is not utilized by crops.^35,36^ Although not evaluated in this study, the use of a simple test that could be employed by farmers to determine on-site urease activity from soils may help decide appropriate fertilizer types for a given region. The field test may also be useful for screening environmental organisms with high urease activity that can be used for engineering applications. Urease-driven MICP has been explored in recent years for cementation and restoration of diverse structures, from art sculptures to buildings, as well as bedrock plugging for enhanced oil recovery.^3,13,37^ Rapid on-site evaluation of urease activity could help predict the efficiency of restoration approaches, instead of waiting for long-term reactive solutions.

A simple and economical test to detect enzymatic activity in the field may also help us understand the microbial processes that contribute to the chemistry and mineralogy of poorly studied sites. For example, though calcareous fens and other peatland ecosystems are extensive in some regions and are relevant carbon sinks,^38^ little information exists about the activity of their microbial communities, in particular the activities of urease and CA. We showed here not only that calcareous fen microbes have the potential to express urease, but also that their urease is active *in situ*, where urease-driven MICP could occur. Though the increasing use of fertilizers has been linked to ecosystem damage, it may be possible to encourage the use of urea-based fertilizers in regions that are hydrologically connected to calcium-rich areas (such as calcareous fens) where indigenous microbes are known to drive MICP, resulting in a sustainable carbon sequestration alternative.^39^

An economical test may be useful for educational demonstrations, and also could prove valuable to determine temporal variations of environmental metabolisms along different seasons, days, or even hours, which may otherwise prove difficult because of budget constraints. In-field tests may be convenient to assess the influence of MICP on microbialites, particularly where bicarbonate transport and urease-related genes are known to be present,^40^ which could help us understand the elusive metabolisms involved in ancient microbialite formation.^41,42^

Top-down molecular studies of microbial communities and their environmental effects have exploded in the past decade, increasing our understanding of uncultivable microorganisms as well as the diversity of distinct environments in a microbe-dominated Earth. The information currently obtained via high-throughput sequencing of environmental microbes should be complemented with field activity assays to assess not only metabolic potential, but also microbial activity in a given environment. Enzymatic activities not only represent protein expression, but also the microbial effects on the environment, which in this study has been related to carbonate precipitation. We anticipate that field-based bottom-up approaches will aid in addressing challenging questions, such as determining the microbial role in mineral formation, as well as providing new eco-friendly technologies for engineering challenges. To our knowledge, this is the first field environmental urease and CA evaluation using an inexpensive and fast method designed with conventional laboratory materials. We expect that a variety of other environments can be tested using this method by other researchers, expanding our knowledge of environmental protein expression and its effects on MICP.

## ASSOCIATED CONTENT

### Supporting Information

Preparation protocols for the field urease and CA activity assays, methods for protein extraction and microscope visualization, and related field images and sampling information.

## ACKNOWLEDGMENT

We gratefully thank Michael Rosen, Matt Oberhelman, Kim Lapakko, Beverly Flood, Javier García Barriocanal and Barbara MacGregor for field/laboratory support and helpful discussions. The authors declare no competing financial interest. Parts of this work were carried out in the Characterization Facility, University of Minnesota, which receives partial support from NSF through the MRSEC program. This research was funded by a NASA award NNX14AK20G to JVB. FMF acknowledge the support from the UMN Graduate School DDF, Fulbright #15150776 and CONICYT folio-72160214 fellowships. KH was supported by a MnDRIVE Environment initiative grant at the University of Minnesota.

## REFERENCES

(1) Sarayu, K.; Iyer, N. R.; Murthy, A. R. Exploration on the Biotechnological Aspect of the Ureolytic Bacteria for the Production of the Cementitious Materials--a Review. Appl. Biochem. Biotechnol. 2014, 172 (5), 2308–2323. https://doi.org/10.1007/s12010-013-0686-0.

(2) Bose, H.; Satyanarayana, T. Microbial Carbonic Anhydrases in Biomimetic Carbon Sequestration for Mitigating Global Warming: Prospects and Perspectives. Front. Microbiol. 2017, 8, 1615. https://doi.org/10.3389/fmicb.2017.01615.

(3) Krajewska, B. Urease-Aided Calcium Carbonate Mineralization for Engineering Applications: A Review. J. Adv. Res. 2018, 13, 59–67. https://doi.org/10.1016/j.jare.2017.10.009.

(4) Seifan, M.; Berenjian, A. Microbially Induced Calcium Carbonate Precipitation: A Widespread Phenomenon in the Biological World. Appl. Microbiol. Biotechnol. 2019, 103 (12), 4693–4708. https://doi.org/10.1007/s00253-019-09861-5.

(5) Zhu, T.; Dittrich, M. Carbonate Precipitation through Microbial Activities in Natural Environment, and Their Potential in Biotechnology: A Review. Front. Bioeng. Biotechnol. 2016, 4. https://doi.org/10.3389/fbioe.2016.00004.

(6) Krausfeldt, L. E.; Farmer, A. T.; Castro Gonzalez, H. F.; Zepernick, B. N.; Campagna, S. R.; Wilhelm, S. W. Urea Is Both a Carbon and Nitrogen Source for Microcystis Aeruginosa: Tracking ^13^C Incorporation at Bloom PH Conditions. Front. Microbiol. 2019, 10, 1064. https://doi.org/10.3389/fmicb.2019.01064.

(7) Kumar, R. S. S.; Ferry, J. G. Prokaryotic Carbonic Anhydrases of Earth’s Environment. Subcell. Biochem. 2014, 75, 77–87. https://doi.org/10.1007/978-94-007-7359-2_5.

(8) Bachmeier, K. L.; Williams, A. E.; Warmington, J. R.; Bang, S. S. Urease Activity in Microbiologically-Induced Calcite Precipitation. J. Biotechnol. 2002, 93 (2), 171–181. https://doi.org/10.1016/s0168-1656(01)00393-5.

(9) Okwadha, G. D. O.; Li, J. Optimum Conditions for Microbial Carbonate Precipitation. Chemosphere 2010, 81 (9), 1143–1148. https://doi.org/10.1016/j.chemosphere.2010.09.066.

(10) Achal, V.; Pan, X. Characterization of Urease and Carbonic Anhydrase Producing Bacteria and Their Role in Calcite Precipitation. Curr. Microbiol. 2011, 62 (3), 894–902. https://doi.org/10.1007/s00284-010-9801-4.

(11) Fujita, Y.; Taylor, J. L.; Gresham, T. L. T.; Delwiche, M. E.; Colwell, F. S.; Mcling, T. L.; Petzke, L. M.; Smith, R. W. Stimulation of Microbial Urea Hydrolysis in Groundwater to Enhance Calcite Precipitation. Environ. Sci. Technol. 2008, 42 (8), 3025–3032. https://doi.org/10.1021/es702643g.

(12) Cuthbert, M. O.; McMillan, L. A.; Handley-Sidhu, S.; Riley, M. S.; Tobler, D. J.; Phoenix, V. R. A Field and Modeling Study of Fractured Rock Permeability Reduction Using Microbially Induced Calcite Precipitation. Environ. Sci. Technol. 2013, 47 (23), 13637–13643. https://doi.org/10.1021/es402601g.

(13) Dhami, N. K.; Reddy, M. S.; Mukherjee, A. Biomineralization of Calcium Carbonates and Their Engineered Applications: A Review. Front. Microbiol. 2013, 4, 314. https://doi.org/10.3389/fmicb.2013.00314.

(14) Gat, D.; Ronen, Z.; Tsesarsky, M. Soil Bacteria Population Dynamics Following Stimulation for Ureolytic Microbial-Induced CaCO_3_ Precipitation. Environ. Sci. Technol. 2016, 50 (2), 616–624. https://doi.org/10.1021/acs.est.5b04033.

(15) Graddy, C. M. R.; Gomez, M. G.; Kline, L. M.; Morrill, S. R.; DeJong, J. T.; Nelson, D. C. Diversity of Sporosarcina-like Bacterial Strains Obtained from Meter-Scale Augmented and Stimulated Biocementation Experiments. Environ. Sci. Technol. 2018, 52 (7), 3997–4005. https://doi.org/10.1021/acs.est.7b04271.

(16) Power, I. M.; Harrison, A. L.; Dipple, G. M. Accelerating Mineral Carbonation Using Carbonic Anhydrase. Environ. Sci. Technol. 2016, 50 (5), 2610–2618. https://doi.org/10.1021/acs.est.5b04779.

(17) Mohsenpour, M.; Noormohammadi, Z.; Irani, S.; Amirmozafari, N. Expression of an Environmentally Friendly Enzyme, Engineered Carbonic Anhydrase, in Escherichia coli. Int. J. Environ. Res. 2019, 13, 295–301. https://doi.org/10.1007/s41742-019-00178-9.

(18) Thillainayagam, A. V.; Arvind, A. S.; Cook, R. S.; Harrison, I. G.; Tabaqchali, S.; Farthing, M. J. Diagnostic Efficiency of an Ultrarapid Endoscopy Room Test for Helicobacter Pylori. Gut 1991, 32 (5), 467–469. https://doi.org/10.1136/gut.32.5.467.

(19) Ross, P.; Behar, M. Test Strip for h. Pylori Detection. US20120094371A1, April 19, 2012.

(20) Fehder, C. G. Carbon Dioxide Indicator Device. US4728499A, March 1, 1988.

(21) Dean, W. E.; Gorham, E.; Swaine, D. J. Geochemistry of Surface Sediments of Minnesota Lakes. In Elk Lake, Minnesota: Evidence for Rapid Climate Change in the North-Central United States; Bradbury, J. P., Dean, W. E., Eds.; Geological Society of America Special Paper 276: Boulder, Colorado, 1993; pp 115–133. https://doi.org/10.1130/SPE276-p115.

(22) Almendinger, J. E.; Leete, J. H. Peat Characteristics and Groundwater Geochemistry of Calcareous Fens in the Minnesota River Basin, U.S.A. Biogeochemistry 1998, 43 (1), 25. https://doi.org/10.1023/A:1005905431071.

(23) Hesse, K.-F.; Küppers, H. Refinement of the Structure of Ikaite, CaCO_3_·6H_2_O. Z. Krist.-Cryst. Mater. 1983, 163 (3-4), 227–231. https://doi.org/10.1524/zkri.1983.163.3-4.227.

(24) Mobley, H. L.; Island, M. D.; Hausinger, R. P. Molecular Biology of Microbial Ureases. Microbiol. Rev. 1995, 59 (3), 451–480.

(25) Zhou, Y.; Zhang, X.; Li, X.; Jia, P.; Dai, R. Evaluation of Changes in Microcystis Aeruginosa Growth and Microcystin Production by Urea via Transcriptomic Surveys. Sci. Total Environ. 2019, 655, 181–187. https://doi.org/10.1016/j.scitotenv.2018.11.100.

(26) Paerl, H. W.; Scott, J. T.; McCarthy, M. J.; Newell, S. E.; Gardner, W. S.; Havens, K. E.; Hoffman, D. K.; Wilhelm, S. W.; Wurtsbaugh, W. A. It Takes Two to Tango: When and Where Dual Nutrient (N & P) Reductions Are Needed to Protect Lakes and Downstream Ecosystems. Environ. Sci. Technol. 2016, 50 (20), 10805–10813. https://doi.org/10.1021/acs.est.6b02575.

(27) Glibert, P. M.; Maranger, R.; Sobota, D. J.; Bouwman, L. The Haber Bosch-Harmful Algal Bloom (HB-HAB) Link. Environ. Res. Lett. 2014, 9, 105001. https://doi.org/10.1088/1748-9326/9/10/105001.

(28) Kafarski, P.; Talma, M. Recent Advances in Design of New Urease Inhibitors: A Review. J. Adv. Res. 2018, 13, 101–112. https://doi.org/10.1016/j.jare.2018.01.007.

(29) Amtul, Z.; Rahman, A.-U.; Siddiqui, R. A.; Choudhary, M. I. Chemistry and Mechanism of Urease Inhibition. Curr. Med. Chem. 2002, 9 (14), 1323–1348. https://doi.org/10.2174/0929867023369853.

(30) Zimmerman, S. A.; Ferry, J. G.; Supuran, C. T. Inhibition of the Archaeal Beta-Class (Cab) and Gamma-Class (Cam) Carbonic Anhydrases. Curr. Top. Med. Chem. 2007, 7 (9), 901–908. https://doi.org/10.2174/156802607780636753.

(31) Zimmerman, S.; Innocenti, A.; Casini, A.; Ferry, J. G.; Scozzafava, A.; Supuran, C. T. Carbonic Anhydrase Inhibitors. Inhibition of the Prokariotic Beta and Gamma-Class Enzymes from Archaea with Sulfonamides. Bioorg. Med. Chem. Lett. 2004, 14 (24), 6001–6006. https://doi.org/10.1016/j.bmcl.2004.09.085.

(32) Giri, A.; Banerjee, U. C.; Kumar, M.; Pant, D. Intracellular Carbonic Anhydrase from Citrobacter Freundii and Its Role in Bio-Sequestration. Bioresour. Technol. 2018, 267, 789–792. https://doi.org/10.1016/j.biortech.2018.07.089.

(33) Dhami, N. K.; Reddy, M. S.; Mukherjee, A. Synergistic Role of Bacterial Urease and Carbonic Anhydrase in Carbonate Mineralization. Appl. Biochem. Biotechnol. 2014, 172 (5), 2552–2561. https://doi.org/10.1007/s12010-013-0694-0.

(34) Yoshida, N.; Higashimura, E.; Saeki, Y. Catalytic Biomineralization of Fluorescent Calcite by the Thermophilic Bacterium Geobacillus Thermoglucosidasius. Appl. Environ. Microbiol. 2010, 76 (21), 7322–7327. https://doi.org/10.1128/AEM.01767-10.

(35) Cantarella, H.; Otto, R.; Soares, J. R.; Silva, A. G. de B. Agronomic Efficiency of NBPT as a Urease Inhibitor: A Review. J. Adv. Res. 2018, 13, 19–27. https://doi.org/10.1016/j.jare.2018.05.008.

(36) Modolo, L. V.; da-Silva, C. J.; Brandão, D. S.; Chaves, I. S. A Minireview on What We Have Learned about Urease Inhibitors of Agricultural Interest since Mid-2000s. J. Adv. Res. 2018, 13, 29–37. https://doi.org/10.1016/j.jare.2018.04.001.

(37) Phillips, A. J.; Cunningham, A. B.; Gerlach, R.; Hiebert, R.; Hwang, C.; Lomans, B. P.; Westrich, J.; Mantilla, C.; Kirksey, J.; Esposito, R.; et al. Fracture Sealing with Microbially-Induced Calcium Carbonate Precipitation: A Field Study. Environ. Sci. Technol. 2016, 50 (7), 4111–4117. https://doi.org/10.1021/acs.est.5b05559.

(38) Lunt, P. H.; Fyfe, R. M.; Tappin, A. D. Role of Recent Climate Change on Carbon Sequestration in Peatland Systems. Sci. Total Environ. 2019, 667, 348–358. https://doi.org/10.1016/j.scitotenv.2019.02.239

(39) Mitchell, A. C.; Dideriksen, K.; Spangler, L. H.; Cunningham, A. B.; Gerlach, R. Microbially Enhanced Carbon Capture and Storage by Mineral-Trapping and Solubility-Trapping. Environ. Sci. Technol. 2010, 44 (13), 5270–5276. https://doi.org/10.1021/es903270w.

(40) Warden, J. G.; Casaburi, G.; Omelon, C. R.; Bennett, P. C.; Breecker, D. O.; Foster, J. S. Characterization of Microbial Mat Microbiomes in the Modern Thrombolite Ecosystem of Lake Clifton, Western Australia Using Shotgun Metagenomics. Front. Microbiol. 2016, 7, 1064. https://doi.org/10.3389/fmicb.2016.01064.

(41) Bosak, T.; Greene, S.; Newman, D. K. A Likely Role for Anoxygenic Photosynthetic Microbes in the Formation of Ancient Stromatolites. Geobiology 2007, 5 (2), 119–126.

(42) Bosak, T.; Newman, D. K. Microbial Nucleation of Calcium Carbonate in the Precambrian. Geology 2003, 31 (7), 577–580. https://doi.org/10.1130/0091-7613(2003)031<0577:MNOCCI>2.0.CO;2.

