## Supporting Information for "Detection of urease and carbonic anhydrase activity using a rapid and economical field test to assess microbially-induced carbonate precipitation"

Number of pages: 23
Number of figures: 8
Number of tables: 1

### SUPPORTING METHODS

#### 1. Preparation of loading and detection strips

#### 1.1. Urease assay protocol

The urease assay is based on a pH-sensitive strip containing phenol red at pH 6.0, which changes its color from yellow to red upon exposure to $NH_3$ fumes from a sample-loading strip containing urea.[1] The preparation of the strips and assembling of the urease tests were performed according to the following procedure:

1.1.1. The urease detection strip is based on a phenol red pH indicator: Add 0.02 g of phenol red (Sigma-Aldrich, St. Louis, Missouri, USA) in 100 mL of distilled water and adjust the pH to 6.0 with diluted HCl. Phenol red has a very low buffer capacity, impending any pH adjustments beyond its $pK_a$ (~8.0); therefore, use a 50 mM (or less) HCl solution when approaching pH 6.

1.1.2. The urease sample loading strip is based on a 0.6 M urea buffer: Add 1.8 g of urea (Thermo Fisher Scientific, Waltham, MA, USA) to 50 mL of a 0.4 mM EDTA (Thermo Fisher Scientific) solution. Chelating agents (such as EDTA) are not necessary to detect the urease activity; however, it may protect urease from inhibitory metals in the sample matrix. To prepare the loading strip with urease inhibitor (negative control), add 0.0086 g of phenyl phosphorodiamidate (PPD; Alfa Aesar, Ward Hill, Massachusetts, USA) to the 50 mL urea solution, yielding a final concentration of 1 mM PPD.

1.1.3. In a flat glass pan, immerse a cellulose fiber sheet grade P8 (Thermo Fisher Scientific, Cat. 09-795D, Waltham, MA, USA) in fresh phenol red buffer or fresh urea buffer to prepare the indicator and the loading strips, respectively. Use no less than 50 mL per 100 cm$^2$ of paper sheet. Rock the pan to impregnate the paper thoroughly for 30 seconds, flip the sheet and repeat for additional 30 seconds.

1.1.4. Using clean tweezers, take the cellulose paper and extended it on a dry flat glass surface. Lean the glass at ~45° to drain excess solution from the paper and leave at room temperature for ~10 h until it dries completely.

1.1.5. Cut 6.5 mm diameter circles using a hole punch cutter, or alternatively cut strips of ~4 × 8 mm from the dried indicator and loading papers.

1.1.6. To assemble the test, place an indicator paper circle inside the cap of a Seal-Rite 0.5-mL microcentrifuge tube (USA Scientific, Inc., Ocala, Florida, USA), so the paper is visible once the tube is closed (see illustrations in **Figure 1**). Put the loading paper inside the tube. The urease test, prepared as described here in closed microcentrifuge tubes, lasts at least one year stored at room temperature and protected from light with no decrease in sensitivity. The negative control containing PPD, however, expires after approximately two to three months stored at room temperature, because the PPD inhibitor is sensitive to oxygen. After three months, the PPD-impregnated strip is unable to inhibit high urease concentrations and may result in a false positive reaction, although less intense than the urease test without inhibitor. We recommend preparing the inhibitor-impregnated loading strips from a fresh PPD solution a few days prior to use, or alternatively, storing the loading strips under anoxic conditions and protected from light.

1.1.7. Evaluate urease activity by placing a wet biofilm sample or extract in direct contact with the sample-loading strip and close the tube. Incubate at room temperature for 5–60 minutes. Longer incubation times may be useful to detect low urease levels, however, the presence of urea may induce urease expression during long incubations, resulting in a signal that may not reflect in-field microbial conditions. During the test, it is also important to prevent any contact of the sample with the indicator strip to avoid a color change derived from alkaline or acidic samples that may turn the indicator. The dry format of the test yields a faster reaction compared to other liquid- or gel-based assays[2,3] and makes it conducive to field deployment.

**1.2. Carbonic anhydrase (CA) assay protocol**

The CA test is based on a pH-sensitive indicator strip of metacresol purple that changes its color in the presence of volatilized $CO_2$ from a bicarbonate solution.[4] Other pH indicators may be used as well. We additionally tested thymol blue and phenol red, however, metacresol purple showed a better color change under the conditions tested. The preparation of the indicator strip and assembling of the CA tests were performed according to the following directions:

1.2.1. Add 0.05 g of metacresol purple (Sigma-Aldrich, St. Louis, Missouri, USA) and 0.34 g of anhydrous $Na_2CO_3$ (Thermo Fisher Scientific, Waltham, MA, USA) to 50 mL of $CO_2$-free ($N_2$-purged) distilled water. Dilute the metacresol purple-carbonate solution by adding 2.5 mL of fresh solution to ~10 mL of $CO_2$-free distilled water and 12.5 mL of glycerol. Dilute the metacresol purple to a concentration of 0.01% by adding $CO_2$-free water to complete 25 mL.

1.2.2. Use 25 mL of freshly diluted (0.01 %) metacresol purple solution to impregnate a cellulose fiber paper grade P8 (~50 $cm^2$; Thermo Fisher Scientific, Cat. 09-795D, Waltham, MA, USA) in a small glass container. Mix thoroughly for 30 seconds per side.

1.2.3. Remove the cellulose paper from the metacresol purple solution, quickly drain the excess buffer from the paper and extend it on a flat glass surface. Dry the paper immediately with hot air (using a small heat gun or a hair dryer) within ~15 minutes. The freshly impregnated paper has a slight yellow coloration, which turns purple after complete drying.

1.2.4. Using a hole punch cutter, obtain circles of 6.5 mm diameter (or cut strips of ~4 × 8 mm) from the dried indicator paper. The indicator paper is sensitive to atmospheric $CO_2$ and will turn yellow after 2–3 days of air exposure. Keep the strips in a $CO_2$- and humidity-free atmosphere if not used within two days. We found that the indicator strips last for at least 3 months when stored in a tube containing $CO_2$-sequestering agents, such as calcium hydroxide.

1.2.5. Place an indicator strip inside the cap of a Seal-Rite 0.5-mL microcentrifuge tube (USA Scientific, Inc., Ocala, Florida, USA; see illustrations in **Figure 2**). Immediately before the assay, add 80 μL of a cold 1 M $NaHCO_3$ (Thermo Fisher Scientific) solution pH ~8.3 (0.84 g $NaHCO_3$ in 10 mL distilled water) to the microcentrifuge tube. Close the tube only once the sample (microbial biofilm or extract) has been added and mixed with the bicarbonate solution. Incubate the assay on ice (0–4 °C) and compare the color change (purple to yellow) rate with a negative control assayed simultaneously. The negative control may consist of an assay without a sample (blank) or an assay containing 0.06 mM acetazolamide (Sigma-Aldrich, St. Louis, Missouri, USA) in the bicarbonate solution (by adding 5 μL of a fresh 1 mM acetazolamide solution to the 80 μL 1 M $NaHCO_3$). It is also important to use cold solutions and incubate the assay tubes on ice. Cold incubation, as described in the Wilbur and Anderson method of CA activity quantification,[5] limits the non-enzymatic $CO_2$ hydration and is essential to visualize a reaction time difference between negative controls and CA-incubated reactions.

### 2. Microscope examination

Biomass portions obtained in the field were kept at 4–8 °C for microscopic observation using an Olympus BX61 (Olympus Corporation, Tokyo, Japan) fluorescence microscope after DAPI (4',6-diamidino-2-phenylindole) staining. Fluorescent red light (584–664 nm) emission was used to observe chlorophyll autofluorescence and detect potential photosynthetic organisms.

### 3. Microbial protein extraction

Approximately 4 mL of wet biomass was lysed in the field using a modification of the method for obtaining environmental proteins described by Ogunseitan,[6] which has shown the highest protein recovery and stability compared to other commonly used environmental microbial lysis methods.[7] The wet biomass was mixed in a 15 mL centrifuge tube with 2 mL 50 mM Tris, 10% sucrose, 4 mM EDTA, 1.2 mg lysozyme and a protease inhibitor cocktail (Thermo Fisher Scientific, Cat. 87786, Waltham, MA, USA), adjusted to pH 8.0. The tube was incubated for 1 h on ice and then 2 mL of a solution containing 50 mM Tris pH 8.0, 10% sucrose, 4 mM EDTA, 1% Triton X-100 and 1× protease inhibitor cocktail was added, mixed with the previous solution and immediately frozen in dry ice for transportation to the laboratory. The frozen aliquot was then subjected to 4 freeze-thaw cycles (-80 °C to 25 °C) and centrifuged at 7,000 × g and 4 °C for 1 h. A total of 10 mM $MgSO_4$ and 2.5 U DNase I (Thermo Fisher Scientific, Cat. 89836) were added to the lysate supernatant and incubated in ice for 1 h with occasional, gently mixing. A Bradford assay was used for protein quantification following the manufacture's instructions (Bio-Rad Laboratories Inc., Hercules, California) and the lysates were then used for urease and CA activity assays.

**4. Urease and CA activity assays of extracts**

CA activity was evaluated using a modification of the Wilbur & Anderson method.[5] Ten micrograms of protein in 6 mL of cold 0.02 M Tris buffer adjusted to pH 8.0 with $H_2SO_4$ was agitated in a magnetic stirrer at 700 rpm (Thermo Fisher Scientific, Fisherbrand$^{TM}$ Isotemp$^{TM}$) and kept at 0–4 °C. The enzymatic reaction was initiated by introducing 4 mL of cold $CO_2$-saturated water using a five-mL syringe. The time for the pH drop from 8.3 to 6.3 was recorded and compared to a blank assay without protein lysate. One unit of CA activity was defined as $2(t_0\text{-}t)/t$, where t is the time for the pH drop from 8.3 to 6.3 in the presence of the lysate and $t_0$ is the same time recorded by the blank.

Urease activity was assayed in microplates (Thermo Fisher Scientific, Corning$^{TM}$ Cat. 07-200-98) using a modification of the phenol-hypochlorite method described by Achal *et al.*[8] Ten micrograms of protein extract were mixed with 0.2 mM Urea in 50 μL of phosphate buffer (100 mM $Na_2HPO_4$ pH 7.0, 200 mM NaCl) and incubated for 20 min at 25 °C. The ammonia released was determined by adding 100 μL phenol nitroprusside solution and 100 μL alkaline hyphochlorite solution (Sigma-Aldrich, St. Louis, Missouri, USA), incubated for 30 min at 37 °C and quantified at 630 nm relative to a control without urea and compared to an ammonium chloride standard curve. One micromolar unit (μU) is defined as the quantity of enzyme that will liberate 1 μmole of ammonia from urea per minute at pH 7.0 at 25 °C.

In-gel enzymatic activity assays were performed following the protonography method described for CA[9] in non-reducing SDS-PAGE, using 100 mM Tris pH 8.5, 0.1 % metacresol purple as a pH indicator solution. The in-gel activity or urease was assayed similarly, using a 100 mM Tris pH 6.5, 0.1 % phenol red as indicator and a 4 % urea, 0.4 mM EDTA solution as substrate.

### SUPPORTING TABLE

**Table S1.** Description of the environmental samples obtained in this study.

| Sample ID | Sample Name | Site Name | Coordinates | Description | Site reference |
|---|---|---|---|---|---|
| 01 | Salt Lake cyanobacterial bloom | Salt Lake, East shore | 44°57'39.7" N 96°26'12.4" W | Bright green surficial biomass suspended at the lake shore. | (10) |
| 02 | Salt Lake filaments | Salt Lake, East shore | 44°57'39.7" N 96°26'12.4" W | Submerged dark green filaments attached to vegetation at the lake shore. | (10) |
| 03 | Fortier-Sioux filaments | Fortier 8 & Sioux Nation WMA Fen | 44°41'21.6" N 96°27'03" W | Green floating filaments containing abundant air bubbles at shallow inundation ponds. | (11) |
| 04 | Black Dog peat biofilm | Black Dog Lake Fen | 44°47'16.5" N 93°16'35.7" W | Green biofilm intermixed with semi-consolidated peat clays and sand at an eroded bank of a fen stream. | (12) |
| 05 | Nicols Meadow filaments | Nicols Meadow Fen – c | 44°49'18.4" N 93°13'14.6" W | Abundant green filaments at an inundation pond next to Nicols Meadow Fen – c. | (11) |
| 06 | Black Dog North orange stream | Black Dog Lake North Fen | 44°49'09.3" N 93°13'28.7" W | Muddy bright orange precipitates at a spring between Nicols Meadow Fen and Black Dog North Fen. | (11) |
| 07 | St. Mary's Spring orange mat | St. Mary's Spring | 44°58'05.5" N 93°14'05.2" W | Abundant orange precipitates at the emergence of an iron-rich seep. | (13) |
| 08 | St. Mary's Spring green mat | St. Mary's Spring | 44°58'05.5" N 93°14'05.2" W | Green microbial mat beneath abundant orange precipitates. | (13) |
| 09 | Soudan seep streamers | Highway near Soudan, MN | 47°50'30.5" N 92°08'56" W | Green and white filaments in a sulfidic stream with orange precipitates and black sediments. | This study |

### SUPPORTING FIGURES

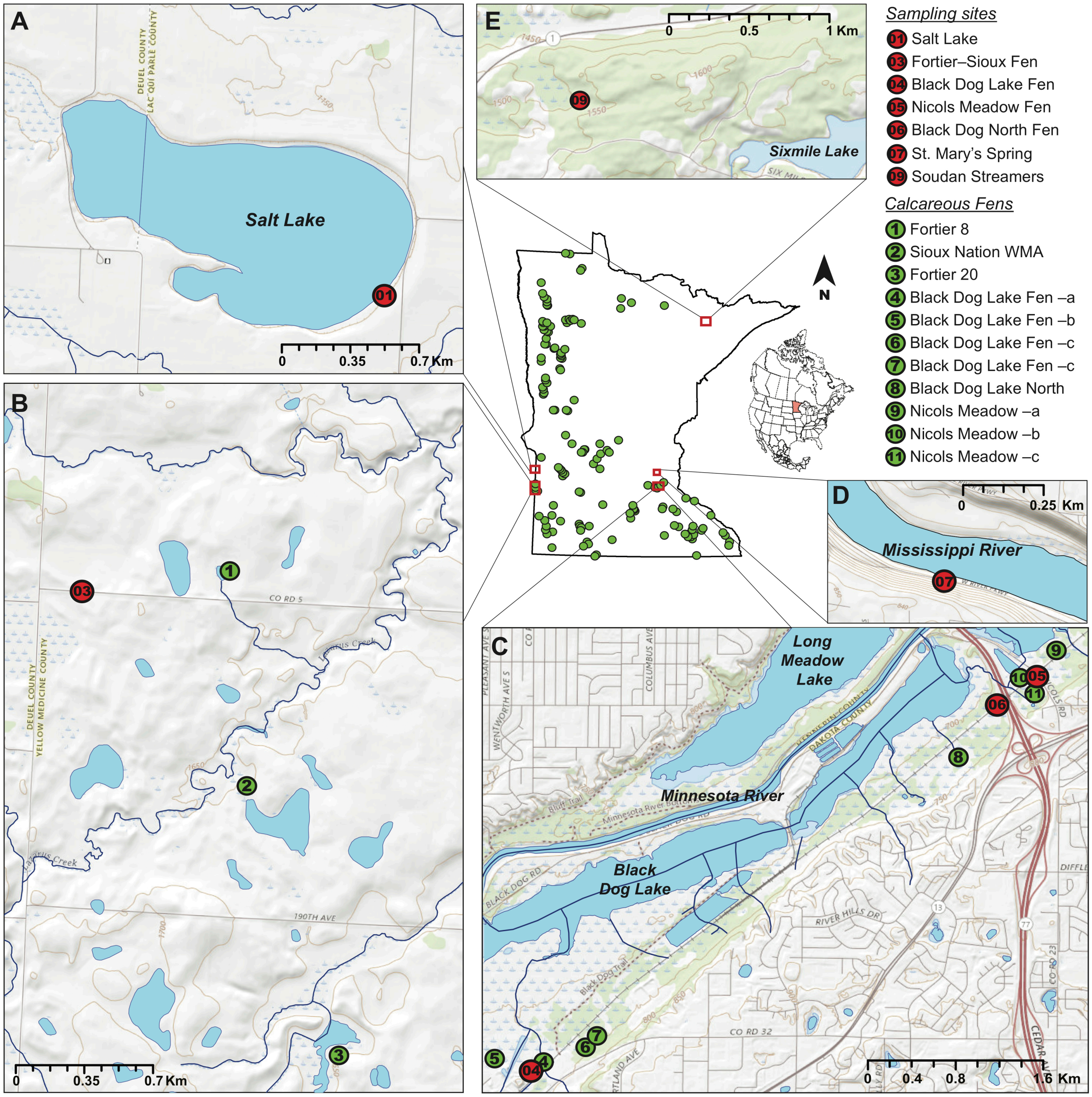


**Figure S1. Microbial sampling sites.** Highlighted contour of Minnesota, USA, indicating with green circles the location of known calcareous fens. Enlarged insets show the sampling locations (red circles) at the Southeastern shore of Salt Lake WMA (A), at a roadside between Fortier 8 and Sioux Nation WMA calcareous fens (B), from a stream at Black Dog Lake Fen, a pond at Nicols Meadow Fen and a creek between Nicols Meadow Fen and Black Dog Lake North Fen

(C), at St. Mary's Springs near the West shore of the Mississippi River (D), and from a sulfidic seep roadside Highway 169, near Soudan, MN (E). Calcareous fen locations and water bodies were obtained from the Minnesota Department of Natural Resources.[12,14] Topographic maps were obtained from the USGS National Geospatial Program, The National Map.[15]

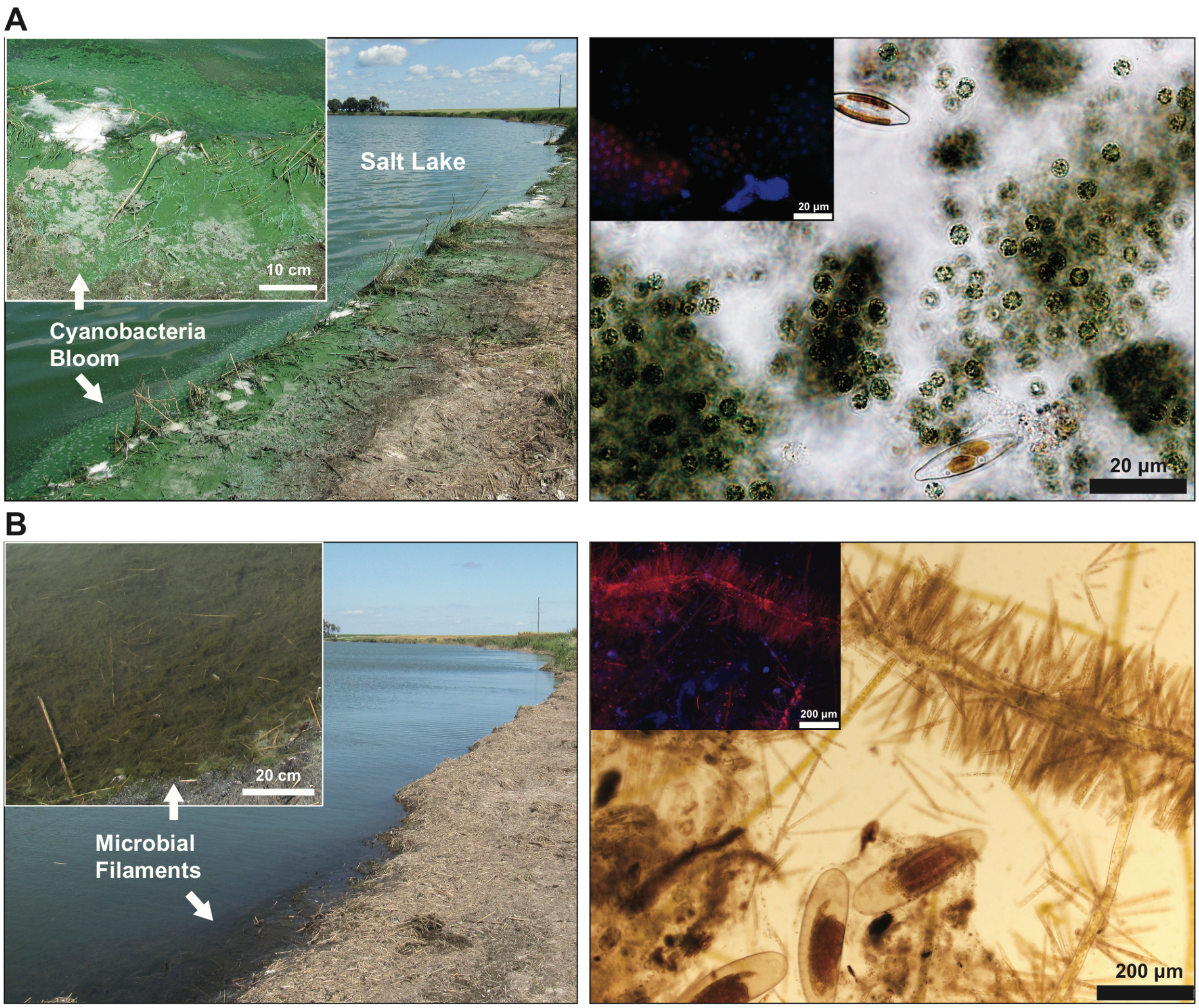


**Figure S2. Microbial biomass in Salt Lake, MN.** Lake shore showing a bright green cyanobacterial bloom (A) sampled in July 2019 and its decline in August 2019 (B), where dark green microbial filaments at the lake shore were sampled instead. Insets show a close-up of the bloom and microfilaments sampled (left panels). Right panels show bright field photomicrographs of the sampled biomass with a merged fluorescence image (insets) of chlorophyll (red) and DAPI (blue) emission. Note that bloom event in A is dominated by a cyanobacterium resembling *Microcystis* spp., whereas the sample taken after the bloom decline is dominated by large green trichome sheaths, probably *Microcoleus* spp., adorned with numerous photosynthetic diatoms.

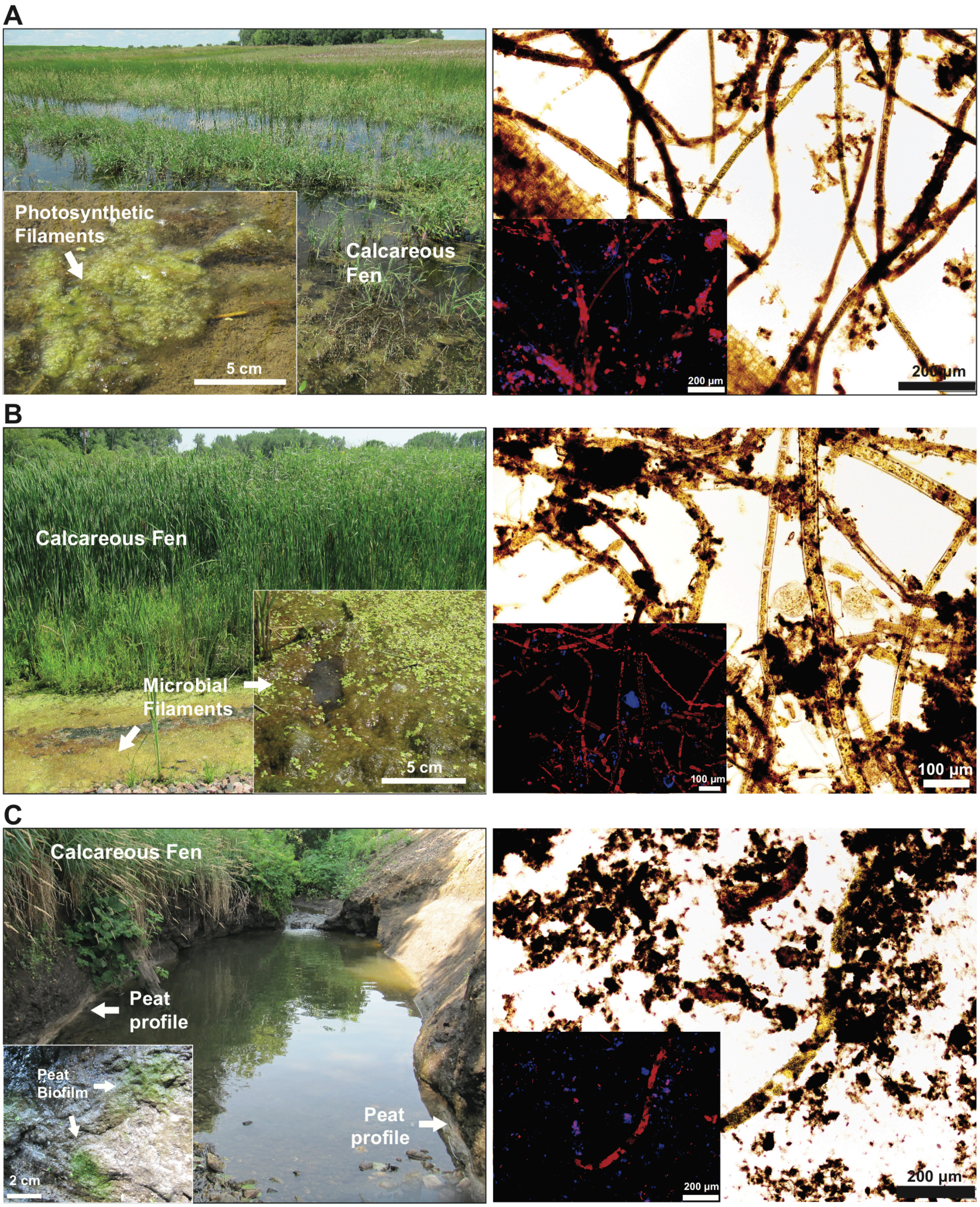


**Figure S3. Microbial filaments at calcareous fens.** Inundation ponds showing green surficial filaments with bubbles at Fortier-Sioux Nation Fens (A) and abundant filaments mixed with aquatic plants (*Spirodela* spp.) at Nicols Meadow fen (B). Green biofilms also colonize the peat exposed in a stream-eroded cliff at Black Dog Lake Fen (C). Insets show a close-up of the biomass sampled (left panels). Right panels show bright field photomicrographs for each sample, with insets of merged fluorescence images of chlorophyll red emission and blue fluorescence from DAPI.

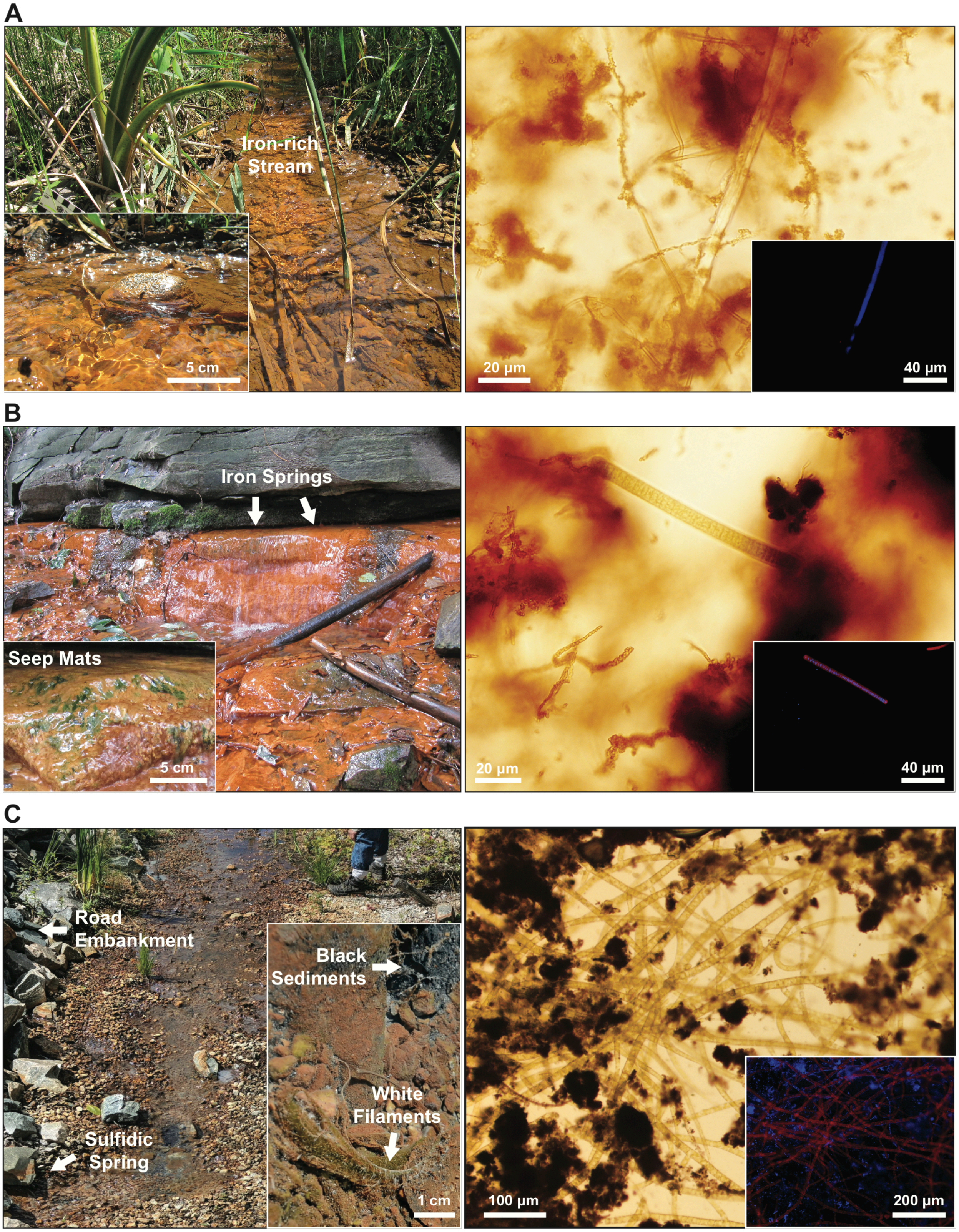
A
Iron-rich
Stream
5 cm
20 µm
40 µm
B
Iron Springs
Seep Mats
5 cm
20 µm
40 µm
C
Road
Embankment
Black
Sediments
White
Filaments
Sulfidic
Spring
1 cm
100 µm
200 µm

188

**Figure S4. Microbial mats at ferrous seeps.** Orange precipitates from a stream between Nicols Meadow Fen and Black Dog Lake North Fen (A), from St. Mary's Spring ferrous seeps (B), and from a sulfidic seep near Soudan, MN (C, field photo credit to Tanner Barnharst). Insets show a close-up of the orange precipitates and microbial mats sampled. Right panels show bright field and fluorescence microscopy (insets) of chlorophyll (red) and DAPI (blue) emission. Note the stalks of iron-oxidizing bacteria in A and B, and abundant cyanobacterial filaments in B and C.

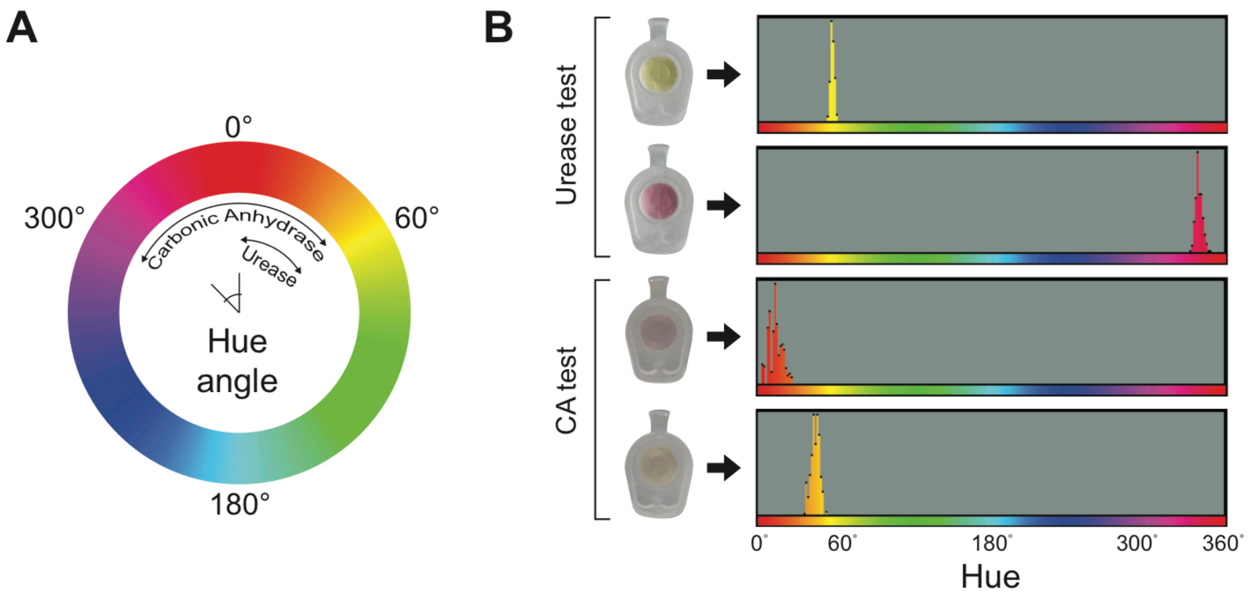


**Figure S5. Quantification of hue values.** (A) Color wheel diagram showing the hue range of the carbonic anhydrase and urease tests. (B) Histograms of hue pixels for representative indicator strip (test tube caps) assays during the enzymatic reaction. The hue was expressed as the average value from assay photos over time.

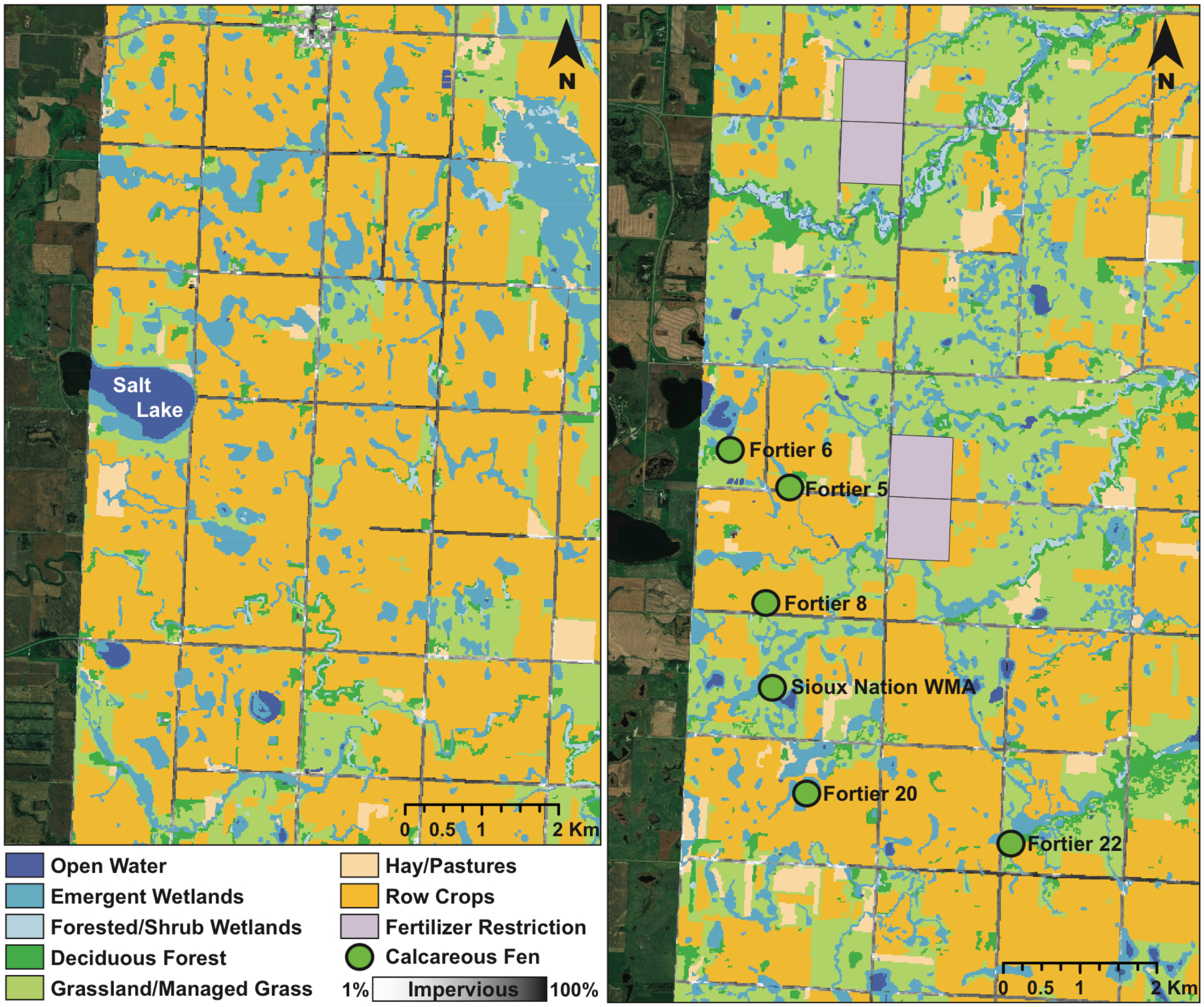


**Figure S6. Land use near Salt Lake and Sioux Nation WMA Fen.** Minnesota land cover classification maps around Salt Lake (A) and Sioux Nation-Fortier fens (B). Classifications according to 2013 Landsat and Lidar data.[16] Note the abundance of agriculture areas near Salt Lake compared to the calcareous fens. Quadrants with nitrogen fertilizer (anhydrous ammonia and urea) application restrictions are based on vulnerable groundwater areas according to the Minnesota Department of Agriculture.[17] Calcareous fen locations were obtained from the Minnesota Department of Natural Resources.[12] South Dakota areas are represented by USGS satellite images.[18]

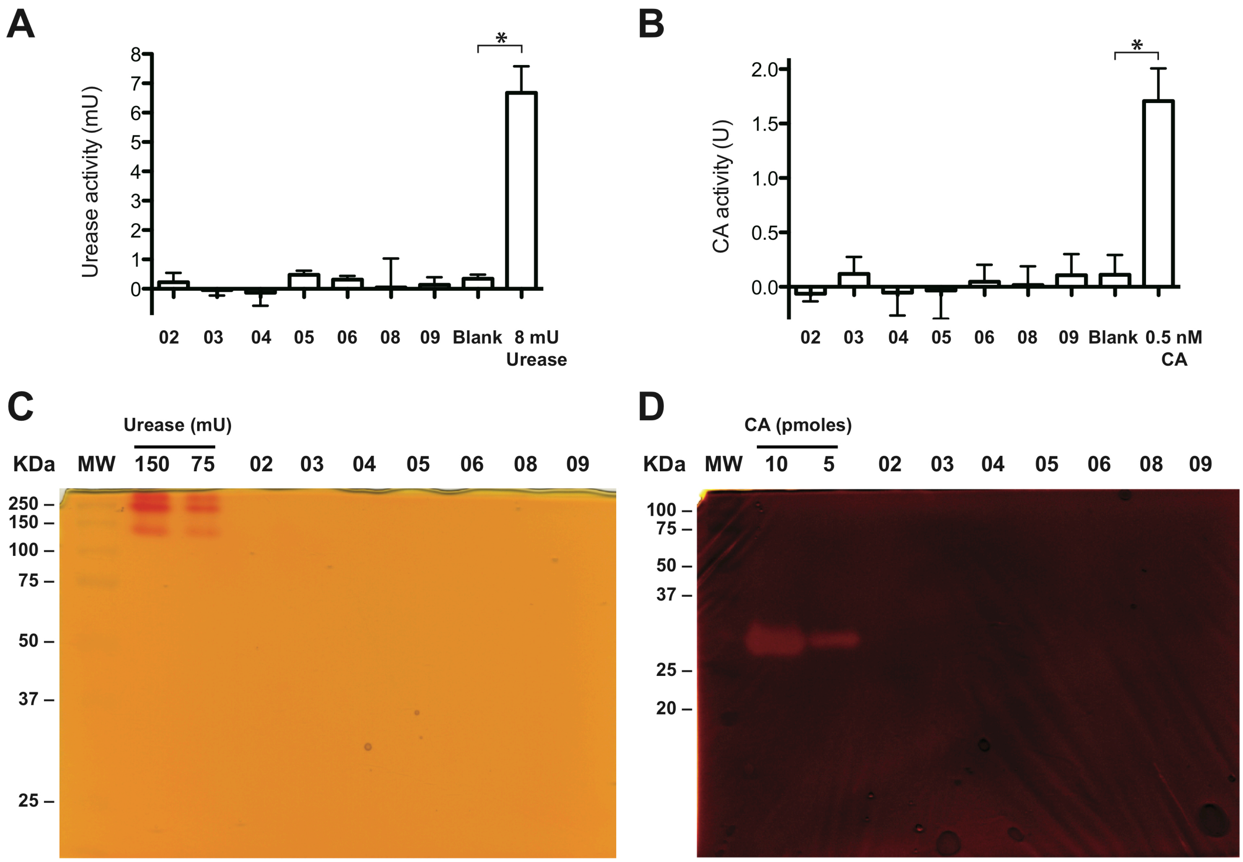


**Figure S7. Urease and carbonic anhydrase activity from protein extracts.** Urease activity using the Berthelot reaction method[8] (A) and CA activity after the Wilbur-Anderson assay[5] (B). A total of 1 µg of protein extracts from Salt Lake filaments (02), Fortier-Sioux filaments (03), Black Dog peat green biofilm (04), Nicols Meadow filaments (05), Black Dog Lake North orange precipitates (06), mixed orange and green mats from St. Mary's Spring (08), and Soudan streamers (09) were used. None of the extracts were significantly higher than a blank assay (*$p < 0.01$, Welch's t-test), which is also observed using in-gel activity assays (C & D). A red band on a yellow background indicates urease activity (C), whereas a yellow band on a purple background indicates CA activity (D).

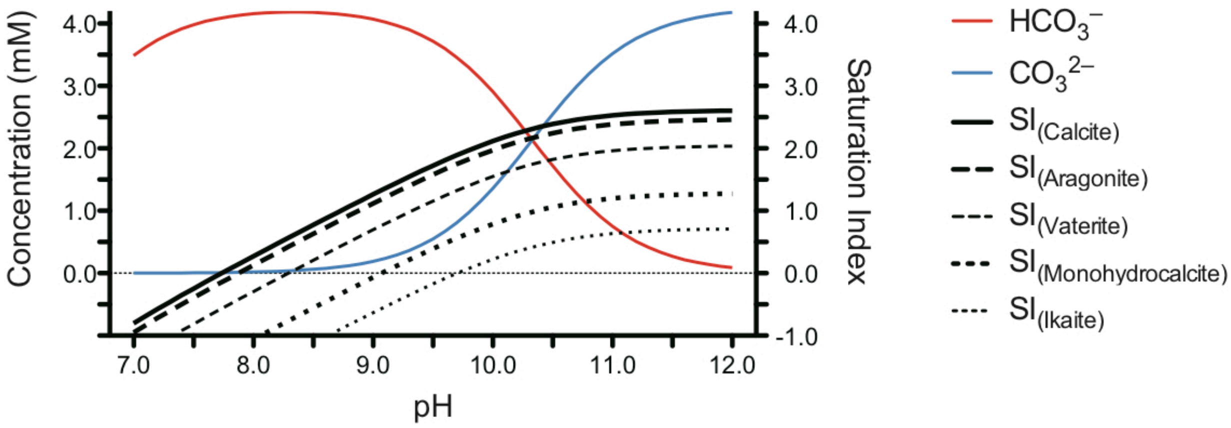


**Figure S8. Salt Lake carbonate speciation and carbonate mineral solubilities.** Bicarbonate and carbonate concentration as a function of pH, considering the alkalinity (234 mg/L as $CaCO_3$) and calcium concentration (16 mM) of Salt Lake. Assuming constant calcium and carbon concentrations, the saturation index (SI) for calcium carbonate minerals increases linearly from pH 9.0 (lake water pH) to 9.6 (pH at the end of microbial incubations without urease inhibitors). Note that above pH 9.6, all metastable calcium carbonate phases are saturated. Solubility values at 25 °C were obtained from Plummer and Busenberg,[19] Kralj and Brečević,[20] and Bischoff *et al.*[21]

### REFERENCES


(1) Ross, P.; Behar, M. Test Strip for h. Pylori Detection. US20120094371A1, April 19, 2012.

(2) Tseng, C.-A.; Wang, W.-M.; Wu, D.-C. Comparison of the Clinical Feasibility of Three Rapid Urease Tests in the Diagnosis of Helicobacter Pylori Infection. *Dig. Dis. Sci.* **2005**, 50 (3), 449–452. https://doi.org/10.1007/s10620-005-2456-5.

(3) Dechant, F.-X.; Dechant, R.; Kandulski, A.; Selgrad, M.; Weber, F.; Reischl, U.; Wilczek, W.; Mueller, M.; Weigand, K. Accuracy of Different Rapid Urease Tests in Comparison with Histopathology in Patients with Endoscopic Signs of Gastritis. *Digestion* **2019**, 1–7. https://doi.org/10.1159/000497810.

(4) Fehder, C. G. Carbon Dioxide Indicator Device. US4728499A, March 1, 1988.

(5) Wilbur, K. M.; Anderson, N. G. Electrometric and Colorimetric Determination of Carbonic Anhydrase. *J. Biol. Chem.* **1948**, 176 (1), 147–154.

(6) Ogunseitan, O. A. Direct Extraction of Proteins from Environmental Samples. *J. Microbiol. Methods* **1993**, 17 (4), 273–281. https://doi.org/10.1016/0167-7012(93)90056-N.

(7) Singleton, I.; Merrington, G.; Colvan, S.; Delahunty, J. S. The Potential of Soil Protein-Based Methods to Indicate Metal Contamination. *Appl. Soil Ecol.* **2003**, 23 (1), 25–32. https://doi.org/10.1016/S0929-1393(03)00004-0.

(8) Achal, V.; Mukherjee, A.; Basu, P. C.; Reddy, M. S. Lactose Mother Liquor as an Alternative Nutrient Source for Microbial Concrete Production by Sporosarcina Pasteurii. *J. Ind. Microbiol. Biotechnol.* **2009**, 36 (3), 433–438. https://doi.org/10.1007/s10295-008-0514-7.

(9) De Luca, V.; Del Prete, S.; Supuran, C. T.; Capasso, C. Protonography, a New Technique for the Analysis of Carbonic Anhydrase Activity. *J. Enzyme Inhib. Med. Chem.* **2015**, 30 (2), 277–282. https://doi.org/10.3109/14756366.2014.917085.

(10) Dean, W. E.; Gorham, E.; Swaine, D. J. Geochemistry of Surface Sediments of Minnesota Lakes. In *Elk Lake, Minnesota: Evidence for Rapid Climate Change in the North-Central United States*; Bradbury, J. P., Dean, W. E., Eds.; Geological Society of America Special Paper 276: Boulder, Colorado, 1993; pp 115–133. https://doi.org/10.1130/SPE276-p115.

(11) Almendinger, J. E.; Leete, J. H. Peat Characteristics and Groundwater Geochemistry of Calcareous Fens in the Minnesota River Basin, U.S.A. *Biogeochemistry* **1998**, 43 (1), 25. https://doi.org/10.1023/A:1005905431071.

(12) Minnesota Department of Natural Resources. Calcareous Fen inventory, 2017, available at https://gisdata.mn.gov/dataset/biota-nhis-calcareous-fens. DNR Hydrography Dataset available at https://gisdata.mn.gov/dataset/water-dnr-hydrography.

(13) Anderson, J. R.; Runkel, A. C.; Tipping, R. G.; Barr, K. D. L.; Alexander, E. C., Jr. Hydrostratigraphy of a Fractured, Urban Aquitard. In *Archean to Anthropocene: Field Guides to the Geology of the Mid-Continent of North America*; Miller, J. D., Jr., Hudak, G. J., Wittkop, C., McLaughlin, P. I., Eds.; Geological Society of America Field Guide 24, 2011; pp 457–475. https://doi.org/10.1130/2011.0024(22).

(14) Minnesota Department of Natural Resources. DNR Hydrography Dataset, 2019, available at https://gisdata.mn.gov/dataset/water-dnr-hydrography.

(15) U.S. Geological Survey. USGS National Geospatial Program, The National Map, 2019, available at https://www.usgs.gov/core-science-systems/national-geospatial-program/national-map.

(16) Rampi, L. P.; Knight, J. F.; Bauer, M. Minnesota Land Cover Classification and Impervious Surface Area by Landsat and Lidar: 2013 Update. Retrieved from the Data Repository for the University of Minnesota, 2016. http://doi.org/10.13020/D6JP4S.

(17) Minnesota Department of Agriculture. Fall Nitrogen Fertilizer Application Restrictions, 2019, available at https://www.mda.state.mn.us/chemicals/fertilizers/nutrient-mgmt/nitrogenplan/mitigation/wrpr/wrprpart1/vulnerableareamap.

(18) U.S. Geological Survey. USGS Geospatial Data Sources, 2019. Satellite imagery available online at https://earthexplorer.usgs.gov/.

(19) Plummer, L. N.; Busenberg, E. The Solubilities of Calcite, Aragonite and Vaterite in $CO_2$-$H_2O$ Solutions Between 0 and 90°C, and an Evaluation of the Aqueous Model for the System $CaCO_3$-$CO_2$-$H_2O$. *Geochim. Cosmochim. Ac.* **1982**, 46 (6), 1011–1040.

(20) Kralj, D.; Brečević, L. Dissolution kinetics and solubility of calcium carbonate monohydrate. *Colloids Surf., A* **1995**, 96 (3), 287–293. https://doi.org/10.1016/0927-7757(94)03063-6.

(21) Bischoff, J. L.; Fitzpatrick, J. A.; Rosenbauer, R. J. The Solubility and Stabilization of Ikaite ($CaCO_3 \cdot 6H_2O$) from 0° to 25°C: Environmental and Paleoclimatic Implications for Thinolite Tufa. *J. Geol.* **1993**, 101 (1), 21–33. https://doi.org/10.1086/648194.
